# The Injury Response to DNA Damage Promotes Anti-Tumor Immunity

**DOI:** 10.1101/2020.04.26.062216

**Authors:** Ganapathy Sriram, Lauren Milling, Jung-Kuei Chen, Wuhbet Abraham, Erika D. Handly, Darrell J. Irvine, Michael B. Yaffe

## Abstract

Inhibition of immune checkpoints has shown promising results in the treatment of certain tumor types. However, the majority of cancers do not respond to immune checkpoint inhibition (ICI) treatment, indicating the need to identify additional modalities that enhance the response to immune checkpoint blockade. In this study, we identified a tumor-tailored approach using *ex-vivo* DNA damaging chemotherapy-treated tumor cells as a live injured cell adjuvant. Using an optimized *ex vivo* system for dendritic cell-mediated T-cell IFN-γ induction in response to DNA-damaged tumor cells, we identified specific dose-dependent treatments with etoposide and mitoxantrone that markedly enhance IFN-γ production by T-cells. Unexpectedly, the immune-enhancing effects of DNA damage failed to correlate with known markers of immunogenic cell death or with the extent of apoptosis or necroptosis. Furthermore, dead tumor cells alone were not sufficient to promote DC cross-presentation and induce IFN-γ in T-cells. Instead, the enhanced immunogenicity resided in the fraction of injured cells that remained alive, and required signaling through the RIPK1, NF-kB and p38MAPK pathways. Direct *in vivo* translation of these findings was accomplished by intra-tumoral injection of *ex vivo* etoposide-treated tumor cells as an injured cell adjuvant, in combination with systemic anti-PD1/CTLA4 antibodies. This resulted in increased intra-tumoral CD103^+^ dendritic cells and circulating tumor antigen-specific CD8^+^ T-cells, leading to enhanced anti-tumor immune responses and improved survival. The effect was abrogated in BATF3-deficient mice indicating that BATF3^+^ DCs are required for appropriate T-cell stimulation by live but injured DNA-damaged tumor cells. Notably, injection of the free DNA-damaging drug directly into the tumor failed to elicit such an enhanced anti-tumor response as a consequence of simultaneous damage to dendritic cells and T-cells. Finally, the DNA damage induced injured cell adjuvant and systemic ICI combination, but not ICI alone, induced complete tumor regression in a subset of mice who were then able to reject tumor re-challenge, indicating induction of a long-lasting anti-tumor immunological memory by the injured cell adjuvant treatment *in vivo*.

## INTRODUCTION

Cytotoxic chemotherapy remains a mainstay of cancer treatment. It is estimated that >60% of all cancer patients will receive some type of initial treatment beyond surgery that includes DNA damaging drugs and/or anti-microtubule agents, particularly in patients with more advanced stage disease (1). These conventional chemotherapy treatments result in extended survival and/or cure, depending on the specific tumor type, however, toxic side effects, and the development of resistance and subsequent tumor relapse is frequently seen.

Therapeutic manipulation of the immune system has now emerged as an alternative approach to anti-cancer therapy, as a consequence of the development of immune checkpoint inhibitors targeting the PD-1/PD-L1 and CTLA-4 axes (2). Certain tumor types show impressive clinical responses to these agents, particularly melanoma (3), non-small cell lung cancer (4,5), and MSI-high colon cancer (6,7). However, the majority of patients with most common tumor types, including breast cancer (8,9), ovarian cancer (10,11), and MSS colon cancer (12) show much lower response rates. It has been estimated that the overall percentage of all cancer patients who will respond to immune checkpoint inhibitors alone is less than 13% (13).

For many tumor types, immunotherapy has been reserved as a second- or third-line treatment option in patients who have failed prior treatment with cytotoxic agents (14). However, early combination of chemotherapy with immune checkpoint inhibitors as a first line therapeutic modality was recently approved for EGFR, ALK, and ROS negative non-small cell lung cancer (NSCLC) using cisplatin and pembrolizumab (15), and for head and neck squamous cell carcinomas (HNSCC) using platinum agents, 5-FU, and pembrolizumab (16).

Data supporting this approach comes from the KEYNOTE-189 trial, which showed a median progression-free survival of 8.8 months in patients with NSCLC that were treated with a combination of cisplatin or carboplatin, pemetrexed, and pembrolizumab, compared to 4.9 months in patients who were treated with chemotherapy alone. However, over 65% of the patients who received this chemotherapy and immunotherapy combination continued to have progressive disease (15). Similarly, the KEYNOTE-048 trial, performed in patients with recurrent unresectable HNSCC in which the tumor contained greater than 1% of cells staining positively for PD-L1 failed to show any improvement in progression-free survival in patients treated with cisplatin or carboplatin, 5-FU, and pembrolizumab, compared to those treated with the same chemotherapy plus cetuximab, although there was an increase in median overall survival from 10.7 months to 13 months when pembrolizumab was included in the combination (16). Clearly, identifying mechanisms that would enhance response rates to the combination of immune checkpoint blockade and chemotherapy, and prolong the durability of the response, remains an unmet clinical need.

Based on our long-standing interest in cross-talk between the DNA damage response and signaling pathways that mediate tumor cell survival, apoptotic cell death, and innate immune activation (17–22), we investigated whether signaling pathways activated in response to specific types of DNA damaging chemotherapy could enhance subsequent anti-tumor immune responses. While the ability of specific chemotherapeutic compounds to enhance cross presentation of tumor antigens by dendritic cells has been characterized as “immunogenic cell death” (23–26), we found that chemotherapy-induced cell stress signaling in live injured cells, but not the presence of dead cells, was the primary determinant of T-cell immunity. This effect seems to be mediated by RIPK1, p38MAPK and NF-kB signaling in the injured tumor cells. Furthermore, we show that direct intra-tumoral injection of *ex vivo* chemotherapy treated cells as an injured cell adjuvant, in combination with systemic ICI, but not systemic ICI alone, drives anti-tumor immunity and tumor regression in murine melanoma models.

## MATERIALS AND METHODS

### Reagents, Cell lines and mouse strains

Mouse GM-CSF and AnnV-FITC were purchased from Biolegend. IL-4 was purchased from Thermo Fisher Scientific. Anti-CD3 (FITC) (145-2C11), Anti-CD8 (APC) (53-6.7), Anti-IFNγ (PE) (XMG1.2), Anti-CD45 (BUV395)(30-F11), Anti-CD24 (APC) (M1/69), Anti-Ly6C (BV605) (AL-21), Anti-F4/80 (BV711) (BM8), Anti-MHCII (PE-Cy7) (M5/14.15.2), Anti-CD11b (BV786) (M1/70), Anti-CD103 (BV421) (2E7) were purchased from ebioscience or Biolegend. H2-K^b^/SIINFEKL-tetramer (PE-conjugated) was purchased from MBL Life Science. Necrostatin-1 and Z-VAD were purchased from Invivogen. Bay 11-7085 and SB202190 were purchased from Sigma. Doxorubicin, Etoposide, Mitoxantrone, Cisplatin, Paclitaxel, Camptothecin, Irinotecan, 5-FU and cylcophosphamide were purchased from LC labs or Sigma. Oxaliplatin was purchased from Tocris Biosciences. An antibody against ovalbumin was purchased from Abcam (Cat # ab17293). PhosphoRIPK1 (S166) (Cat # 31122S) and RIPK1 (Cat # 3493T) antibodies were purchased from Cell Signaling Technology. Calreticulin antibodies were purchased from Invitrogen (Cat # PA3-900) and Cell Signaling Technology (Cat # 12238T). CellTiter-Glo was purchased from Promega. CountBright absolute counting beads for flow cytometry, ACK lysis buffer, Lipofectamine RNAiMax transfection reagent, and LIVE/DEAD Fixable Aqua Dead Cell Stain kit were purchased from Thermo Fisher Scientific. HMGB1 ELISA kit was purchased from IBL international. CD8+ T-cell isolation kit was from STEM cell technologies. Anti-PD1 (clone RMP1-14) and anti-CTLA4 (clone 9D9) were from BioXcell. Anti-Batf3 antibody was purchased from Abcam (#ab211304).

B16F10 cells and MC-38 cells were obtained from ATCC. B16F10 cells were engineered to stably express ovalbumin (B16-Ova cells), as described previously (27). MC-38 Ova cells were generated by transduction of MC-38 cells with pLVX-Ovalbumin-IRES-hygro, selection of stable expression clones using hygromycin, followed by isolation and expansion of single cell clones. Ovalbumin expression was verified by Western blotting (Fig. S1B). Calreticulin siRNA (silencer select ID # s63272) was purchased from Thermo Fisher Scientific.

C57BL/6J WT, BATF3 (-/-), and OT-1 mice were purchased from Jackson laboratories.

### BMDC generation

Bone marrow was harvested from the femurs and tibias of Taconic C57BL/6 mice. The bone marrow was flushed out after nipping one end, and then centrifuged at 15,000 x g for 15s. Following 1 round of RBC lysis with ACK lysis buffer, cells were filtered through a 100µm filter to remove aggregates, re-suspended at 1 × 10^6^ cells/ml, and cultured on a 10 cm bacterial plate (12 million cells per plate) in Iscove’s Modified Dulbecco’s Medium (IMDM) containing 10% FBS with antibiotics, 20 ng/ml each of GM-CSF and IL-4 and 55 µM of β-mercaptoethanol. After 3 days, 75% of the media was replaced with fresh media containing growth factors. Dendritic cells, which were loosely adherent, were harvested by gentle pipetting on day 6 or 7 and used for the assay.

### In vitro cross presentation assay

B16-Ova or MC-38 Ova cells were treated with various doses of chemotherapeutic drugs for 24 h followed by extensive washing in IMDM (10%FBS, P/S). Subsequently 1 × 10^6^ treated cells were co-cultured with 2.5 × 10^5^ BMDC per well of a 24-well plate for each condition tested. After 24 hours of co-culture, supernatants were removed from each well and the BMDC washed 2-3 times in T-cell media (RPMI containing 10% FBS, 20 mM HEPES, 1mM sodium pyruvate, 55 uM β-mercaptoethanol, 2mM L-glutamine, non-essential amino acids and antibiotics). CD8+ OT-I T-cells isolated from spleens of OT-I mice were then co-cultured with the BMDC at 125,000 T-cells per well to achieve an effector to target ratio of 0.5. Where indicated, BMDC and/or T-cells were also exposed to chemotherapy drugs. After a 12-15h incubation, IFN-γ producing T-cells were identified and quantified by intra-cellular cytokine staining and flow cytometry using a BD LSR II or Fortessa flow cytometer. Cells were first gated for CD3 expression, then re-gated for CD8 and IFNγ expression.

In some experiments B16Ova cells were co-treated with 20 µM of Necrostatin-1 or Z-VAD or 10 µM each of SB202190 or Bay 11-7085 and etoposide or mitoxantrone at the concentrations of 10 or 50 µM for 24 hours prior to performance of the above assay.

### Fractionation of live and dead fractions from chemotherapy-treated cells

B16-Ova cells or MC-38-Ova cells were treated with various doses of chemotherapy as indicated in Fig S2 for 24 hours after which the floating fraction of cells was transferred to a separate tube and washed with PBS (for AnnV/DAPI staining) or IMDM (for co-culture with BMDC). The attached fraction was rinsed 1X with PBS, detached using 5 mM EDTA (in PBS), washed with PBS or IMDM and transferred to a separate tube. Separately, cells treated with chemotherapy for 24h were re-plated at 1 million cells per well of a 24-well plate in 500 ul of IMDM (10%FBS; P/S). Cell-free supernatants were collected after a further 24h. As shown in Fig. S2, staining with AnnV and DAPI of the attached and floating fractions after chemotherapy treatment and fractionation revealed that the attached fraction is predominantly AnnV and DAPI double negative suggesting that the majority of cells in this fraction are live injured cells. On the other hand, the floating fraction (labeled as ‘suspension’ in Fig. S2) consists of cells that predominantly stain positive for AnnV and/or DAPI suggesting that the majority of cells in this fraction are dead cells. Lysate of the total chemotherapy-treated cell mixture was generated by three rounds of freeze-thawing by alternate incubations in liquid nitrogen and a 37 C water bath.

### Measurement of immunogenic cell death markers

For measurement of calreticulin surface exposure, B16-Ova cells were treated for 24 hours with various chemotherapy drugs. All attached and floating cells were harvested and washed in staining buffer (PBS containing 0.5% BSA) and incubated with anti-calreticulin antibodies for 1 hour on ice. Cells were washed once in staining buffer and then incubated with secondary AF488-conjugated secondary antibody for 1 hour at room temperature, washed again, re-suspended in staining buffer and analyzed by flow cytometry.

For HMGB1 measurement in cell culture media, B16-Ova cells were treated for 24 hours with various chemotherapy drugs, media was collected, and floating cells removed by centrifugation at 250 x g for 5 min. Cell-free cell culture media was then analyzed by ELISA for HMGB1 according to the manufacturer’s protocol.

For measurement of ATP levels, cell-free culture media obtained as above was analyzed by CellTiter-Glo according to the manufacturer’s protocol. Values were converted to ATP concentrations using a standard curve generated using pure ATP.

*Calreticulin siRNA experimental method:* B16-Ova cells were transfected with calreticulin or control siRNA (30 nM final concentration) using Lipofectamine RNAiMax according to the manufacturer’s protocol. 48 hours post-transfection, cells were used for the *in vitro* cross-presentation assay.

### Cell death and viability assays

For assessment of cell death, floating and attached cells were harvested after 48 hours of treatment with the indicated chemotherapeutic drugs. Attached cells were detached using 5mM EDTA in PBS. The recovered cells were centrifuged at 250 x g for 5 min, washed once in PBS containing 0.9 mM Ca^2+^ and 0.5 mM Mg^2+^ and then stained with AnnV-FITC for 15 minutes in Annexin binding buffer at room temperature according to the manufacturer’s protocol (Biolegend). Cells were co-stained with DAPI at a final concentration of 1 µg/ml for 2 minutes in Annexin binding buffer, brought to a final volume of 500 ul using PBS containing 0.9 mM Ca^2+^ and 0.5 mM Mg^2+^ and analyzed by flow cytometry.

For assays of survival, 15,000 cells were plated per well in a 96-well plate in 100 µl media with 5 replicates per condition. Wells along the four edges of the plate were not used. Following cell attachment, the indicated drugs were added in an equal volume of media, and incubated for an additional 48 hours. The media was then removed and replaced with 100 ul of fresh media at room temperature. Following a 30 minute incubation at room temperature, 50 µl of CellTiter-Glo reagent was added, followed by 2 minutes of gentle mixing. The plate was incubated at room temperature for an additional 10 minutes. 100 µl of supernatant was transferred to a 96-well white opaque plate and luminescence was read on a Tecan microplate reader. Values were normalized to those of DMSO-treated control cells.

### Mouse studies

B16-Ova cells or MC-38 cells (1 × 10^6^) were implanted subcutaneously in the right flank of 7-8 week old female C57BL/6J WT or BATF3 (-/-) mice. After 11-13 days tumors of ∼16 mm^2^ median cross-sectional area were typically detectable by palpation. Mice with tumors were then binned into groups and injected intra-tumorally once a week for 3 weeks with 30 µl of either PBS, free etoposide to achieve a final concentration of 50 µM in the tumor volume, or 1 × 10^6^ etoposide-treated cells (24 hours of drug treatment followed by extensive washing with PBS). Where indicated, groups also received intra-peritoneal injections of 200 µg each of anti-PD1 (clone RMP1-14, BioXCell) and anti-CTLA4 (clone 9D9, BioXcell) twice a week for three weeks.

To enumerate circulating tumor antigen-specific CD8+ T-cells, mice were bled retro-orbitally after the second intra-tumoral dose of PBS, etoposide, or etoposide-treated tumor cells, and H2-K^b^/SIINFEKL-tetramer positive CD8+ T-cells analyzed by flow cytometry. Briefly, 50 µl of whole blood was collected by retro-orbital bleeding, centrifuged at 250 x g for 5 min, followed by 3 rounds of RBC lysis in 200ul of ACK buffer. Cells were then washed once in Tetramer stain buffer (PBS containing 5mM EDTA, 1% BSA and 50 nM Dasatinib), and stained with PE-conjugated Tetramer for 40 min at RT, followed by co-staining with anti-CD8 for 10 min at 4°C. Cells were then stained with DAPI, washed and re-suspended in tetramer stain buffer for flow cytometry analysis.

*Tumor size measurements:* Cross-sectional area of tumors was measured in mm^2^ using calipers every 2-3 days.

In tumor re-challenge experiments, naïve mice controls or mice who had complete tumor regression and remained tumor free for at least 60 days were subcutaneously injected in the left flank (contra-lateral to the initial tumor) with 0.1 × 10^6^ B16-Ova cells, and tumor development was monitored for another 60 days.

### Immunophenotyping

Phenotypic characterization of immune cell populations was performed by flow cytometry. Briefly, tumors were harvested and mashed through a 70 µM filter. Collected cells were washed in FACS buffer (PBS containing 5mM EDTA and 1% BSA), resuspended, and counted. Five million cells from each sample were stained with fluorophore-conjugated antibodies on ice for 30 min, co-stained with Aqua, washed, re-suspended in 450 µl, supplemented with 50 µl of CountBright absolute counting beads, and analyzed on a BD LSR Fortessa flow cytometer. DCs were scored as CD45+Ly6C-CD24+MHCII+F480-(CD11b+ or CD103+) cells using the gating strategy described in (28).

### Statistics

All statistical analysis of data was performed using GraphPad Prism software. Comparisons of multiple experimental treatments to a single control condition were analyzed by ANOVA followed by Dunnett’s multiple comparisons test. Comparisons between specific treatment groups were analyzed using a Student’s t-test with Bonferroni correction for multiple hypothesis testing. For greater than three specific comparisons, ANOVA followed by Sidak’s multiple comparisons test was used.

## RESULTS

### Etoposide and mitoxantrone-treated tumor cells can induce DC-mediated OT-I T-cell IFN-γ responses in vitro

To identify how tumor cell stress and injury after DNA-damaging chemotherapy could potentially influence anti-tumor immune function, we treated B16F10 melanoma tumor cells expressing ovalbumin (B16-Ova), or MC-38 colon cancer cells expressing ovalbumin (MC38-Ova) with various doses of the common clinically used chemotherapeutic agents doxorubicin, etoposide, mitoxantrone, cisplatin, oxaliplatin, cyclophosphamide, irinotecan, camptothecin, paclitaxel or 5-FU, and examined them for immunogenicity by assaying for their ability to induce dendritic cell-mediated IFN-γ responses in CD8+ T-cells. As diagrammed in Fig. 1A, following drug treatment for 24 hours, the media was exchanged, and the treated cell mixture (including both injured and dead cells) were then co-cultured with primary bone-marrow derived dendritic cells for an additional 24 hours. Purified CD8+ T-cells obtained from the spleens of OT-1 mice were then added to the drug-treated B16-Ova cells/BMDC co-culture and the appearance of IFN-γ+ CD8+ T-cells quantified 12-15 hrs later by intracellular staining and flow cytometry (Fig.1B). CD8+ T-cells of OT-1 mice express a transgenic T-cell receptor recognizing ovalbumin residues 257-264 in the context of H2-K^b^(29,30).

**Fig 1:**
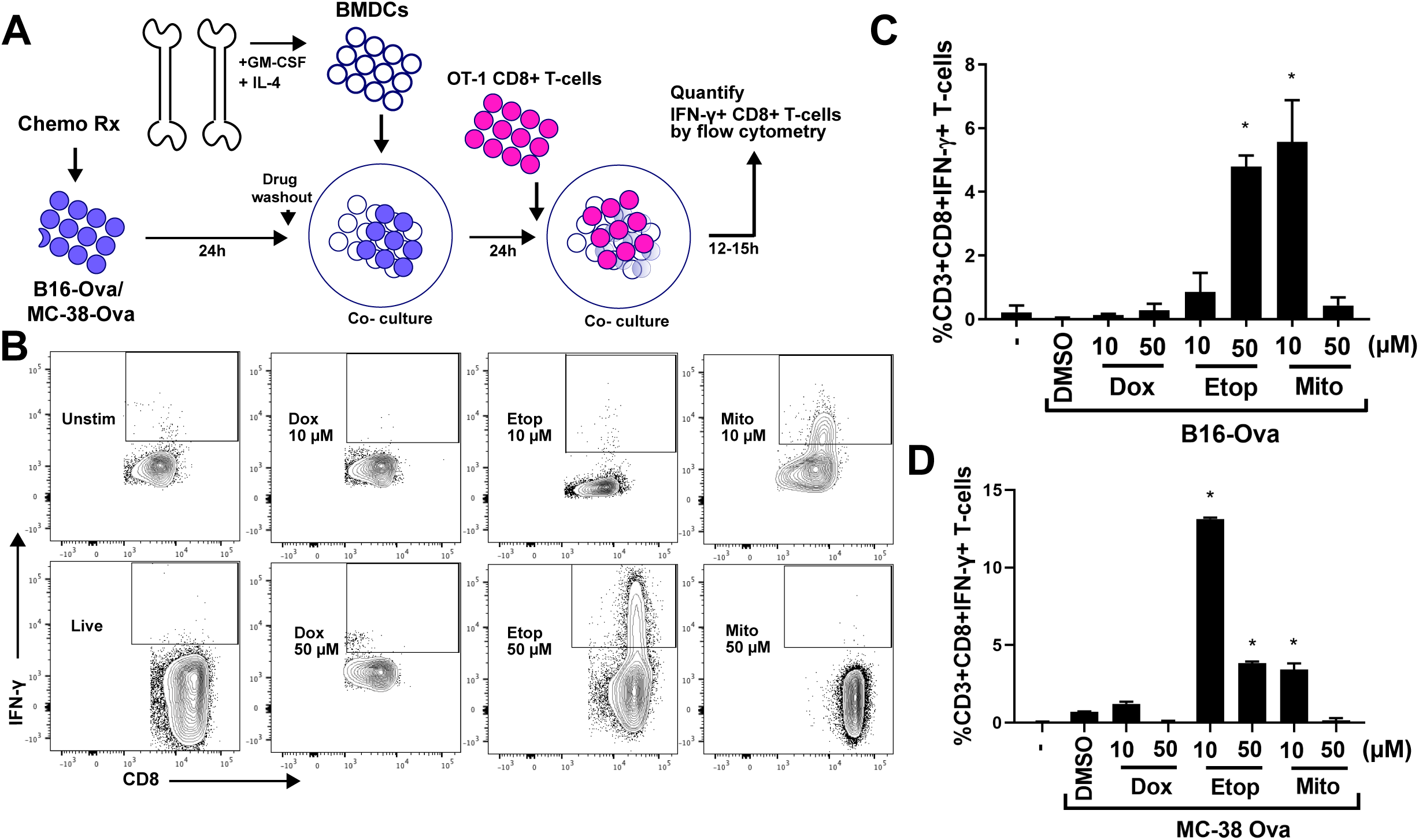
**An experimental system to assess dendritic cell-mediated T-cell IFN-γ responses shows that etoposide- or mitoxantrone-treated B16-Ova or MC-38-Ova cells, when co-cultured with BMDC, effectively induce IFN-γ induction in OT-1 CD8+ T-cells**. **A**. Schematic of the *in vitro* experimental system. B16-Ova or MC-38-Ova cells were treated with DNA-damaging agents for 24 hrs, washed and incubated with primary bone marrow-derived dendritic cells (BMDC) for another 24 hrs. Following this, OT-I CD8^+^ T- cells expressing a TCR transgene that specifically recognizes the Ova-derived peptide SIINFEKL in the context of H2-K^b^ (OT-1)(29,30) were added and evaluated for intracellular IFN-γ 15 hours later. **B**. T-cells were identified by CD3 staining, and re-gated for IFN-γ and CD8 expression. Representative flow cytometry plots showing IFN-γ+ CD8+ T-cell populations (boxed region) induced by treatment of B16-Ova cells with 10µM or 50µM of doxorubicin, etoposide, or mitoxantrone. **C**. Quantification of IFN-γ+ CD8+ T-cells from 5 independent experiments. The first lane (-) indicates the percentage of IFN-γ+ CD8+ T-cells produced by co-culture of BMDCs and T-cells in the absence of B16-Ova cells. Error bars indicate SEM. * indicates p<0.0001 when compared to DMSO-treated control cells using ANOVA followed by Dunnett’s multiple comparisons test. **D**. Quantification of BMDC-mediated induction of IFN-γ+ CD8+ T-cells by chemotherapy-treated MC-38-Ova cells from 3 independent experiments. The first lane (-) indicates the percentage of IFN-γ+ CD8+ T-cells produced by co-culture of BMDCs and T-cells in the absence of MC-38-Ova cells. Error bars represent SEM. * indicates p< 0.0001 when compared to DMSO-treated control using ANOVA followed by Dunnett’s multiple comparisons test.

As shown in Figs. 1C, S1A and 1D, treatment of B16-Ova or MC38-Ova cells with either etoposide or mitoxantrone, was the most effective at inducing DC-mediated IFN-γ in OT-1 CD8+T-cells when the treated cells were co-cultured with BMDC. The effectiveness of these DNA damaging drugs at inducing T-cell IFN-γ responses was highly dose-dependent for each cell line. Surprisingly, in contrast to etoposide and mitoxantrone, another topoisomerase-II inhibitor, doxorubicin, was ineffective at inducing DC-mediated IFN-γ in T-cells, despite causing similar or higher levels of total cell death (Fig. S1A).

### Live injured cells, rather than dead cells, are determinants of DC-mediated IFN-γ induction in T-cells in response to mitoxantrone and etoposide treatment

As shown in Fig. S1C and D, both drugs that effectively induced DC-mediated IFN-γ in CD8+ T-cells induced substantial amounts of apoptotic and non-apoptotic tumor cell death compared to drugs that failed to elicit an immune response, although notably, doxorubicin also caused similar amounts of cell death but was immunologically silent. Curiously, at the doses used in Fig. S1A, the specific doses of mitoxantrone and etoposide that were maximally effective were not the doses that caused the greatest amount of cell death. To investigate if the magnitude of T-cell IFN-γ responses directly correlated with the amount of dead cells present in the treated tumor cell fractions that were co-incubated with BMDC, we treated tumor cells with increasing doses of etoposide or mitoxantrone from 0 to 100 uM. As shown in Fig 2A, B16-Ova cells treated with increasing doses of etoposide, induced a corresponding increase in the magnitude of IFN-γ responses in T-cells (using the assay described in Fig 1A). However, as shown in Fig 2C, the proportion of dead cells (AnnV or DAPI single or double positive) present in the treated tumor cell mixture increases up to ∼ 30% at 25 uM etoposide, but stays unchanged (at ∼ 30%) between 25 and 75 uM and shows only a further small increase (by ∼5%) at 100uM etoposide treatment. On the other hand, B16-Ova cells treated with 5 uM mitoxantrone induced the maximum IFN-γ responses in T-cells among the doses tested, while cells treated with 10 uM mitoxantrone induced a lower IFN-γ response which became undetectable at 25 uM and higher doses (Fig 2B). The dead cell proportion in the mitoxantrone-treated B16-Ova cell mixture is equivalent between 5 and 10 uM (∼ 50%) and increases to greater than 90% at 25uM and higher doses (Fig 2D). Together these results indicate that the proportion of dead cells in both the etoposide and mitoxantrone-treated B16-Ova tumor cell mixtures does not correlate with the magnitude of T-cell IFN-γ responses induced.

**Fig 2:**
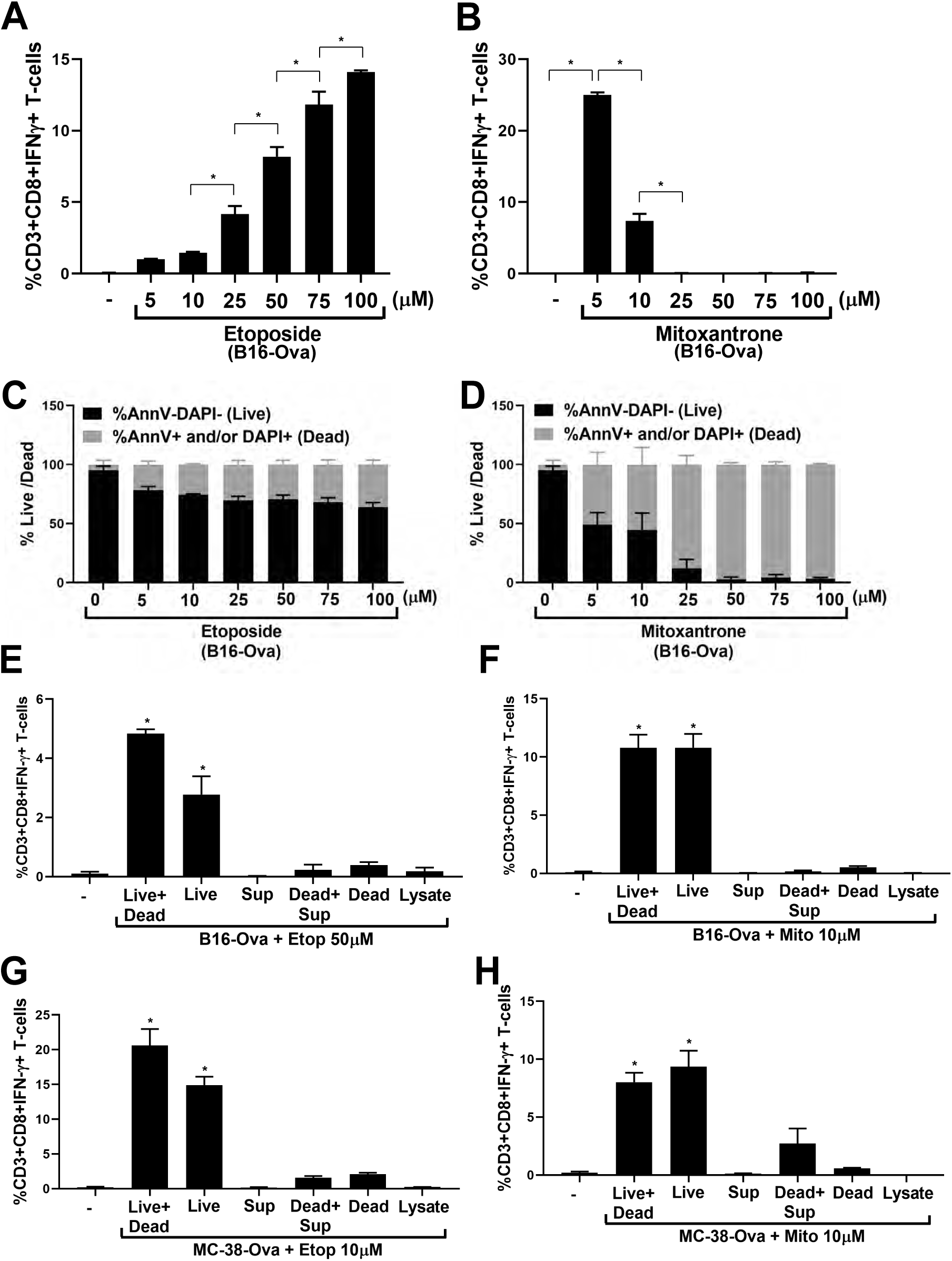
**Live injured cells after treatment by etoposide or mitoxantrone, rather than dead cells, are the primary determinants of DC-mediated T-cell IFN-γ responses** **A**. Quantification (from three independent experiments) of IFN-γ+ CD8+ T-cells induced by co-culture of BMDC with B16-Ova cells treated with etoposide from 0 to 100uM for 24h. The first lane (-) indicates the percentage of IFN-γ+ CD8+ T-cells produced by co-culture of BMDCs and T-cells in the absence of B16-Ova cells. Error bars indicate SEM. *indicates p<0.03 using ANOVA followed by Sidak’s multiple comparisons test. **B**. Quantification (from three independent experiments) of IFN-γ+ CD8+ T-cells induced by co-culture of BMDC with B16-Ova cells treated with mitoxantrone from 0 to 100uM for 24h. The first lane (-) indicates the percentage of IFN-γ+ CD8+ T-cells produced by co-culture of BMDCs and T-cells in the absence of B16-Ova cells. Error bars indicate SEM. *indicates p<0.0001 using ANOVA followed by Sidak’s multiple comparisons test. **C and D**. Quantification (from two to three independent experiments) of the proportion of live (AnnV and DAPI double negative; black bars) and dead (sum total of AnnV and/or DAPI single or double positive; grey bars) cells after treatment of B16-Ova cells for 24h with etoposide or mitoxantrone as indicated. Error bars indicate SEM. **E and F**. Quantification (from three independent experiments) of IFN-γ+ CD8+ T-cells induced by co-culture of BMDC with the indicated B16-Ova cell fractions obtained after treatment with etoposide or mitoxantrone. B16-Ova cells were treated with etoposide at 50 uM or mitoxantrone at 10uM and fractionated into live cells (AnnV and DAPI double negative) and dead cells (AnnV and/or DAPI single or double positive) as described in Methods. Lysate and cell-free supernatants were also obtained as described. BMDC was co-cultured with each of the following fractions or combinations of fractions for 24h before OT-1 CD8+ T-cells were added: (Live+dead) refers to the whole treated cell mixture, (Live) refers to the live cell fraction, (Dead) refers to the dead cell fraction, Sup refers to Cell-free supernatant, (Dead+Sup) refers Dead cells combined with cell-free supernatant, (Dead) refers to the dead cells without cell-free supernatant. Error bars indicate SEM. * indicates p<0.0001 using ANOVA followed by Dunnett’s multiple comparisons test. **G and H**. Quantification (from three independent experiments) of IFN-γ+ CD8+ T-cells induced by co-culture of BMDC with the indicated MC-38-Ova cell fractions obtained after treatment with etoposide or mitoxantrone as described in E and F and in Methods. Error bars indicate SEM. * indicates p<0.0003 using ANOVA followed by Dunnett’s multiple comparisons test.

Since the above results suggested that there was no direct positive correlation between the proportion of dead cells induced by etoposide or mitoxantrone treatment, and the DC-mediated IFN-γ responses in T-cells, we further investigated the specific contribution of the dead and live fractions of tumor cells induced by chemotherapy-treatment. We fractionated the etoposide- and mitoxantrone-treated cell cultures into either cell-free supernatants, supernatants containing dead (AnnV+ and/or DAPI+) cells, or a separate fraction containing only the live (AnnV and DAPI double negative) injured cells (see methods and Fig. S2). As shown in Figs. 2E and F, each fraction was then co-cultured with BMDCs for 24 hrs, followed by the addition of OT-1 CD8+ T-cells for an additional 12-15 hrs, as described above. Neither the cell-free supernatants, nor the supernatants containing the dead cells were capable of inducing DC-mediated T-cell IFN- γ responses. Similarly, lysates generated by subjecting the chemotherapy-treated total cell mixture to three rounds of freeze-thawing (between liquid nitrogen and 37 C), upon co-incubation with BMDC, failed to induce IFN-γ in T-cells. In marked contrast, the fraction containing the adherent live injured cells were the most effective at inducing the expression of IFNγ in OT-1 T-cells. Similar behavior was also noted in the MC-38-Ova cells (Fig 2G and H).

### Conventional markers of immunogenic cell death do not fully explain the T-cell response to mitoxantrone and etoposide treatment

In our *in vitro* assay system, both mitoxantrone and etoposide were found to induce dendritic cell-dependent T-cell IFN-γ responses, in contrast to doxorubicin, which had no effect. Obeid et al (23) reported that mitoxantrone chemotherapy induced strong immunogenic cancer cell death in CT26 mouse colon carcinoma cells, based on the drug’s ability to induce calreticulin exposure on the cell surface. Subcutaneous injection of these drug-treated cells into one flank enabled mice to resist a tumor challenge when live cells were injected into the opposite flank (23). which correlated with resistance to tumor development upon rec-challenge after the drug-treated cells were injected subcutaneously as a prophylactic in a mouse model. In addition to externalized calreticulin, release of HMGB1 and ATP have also been reported to serve as canonical markers of immunogenic cell death (26).

To directly test whether these markers were sufficient to explain DC-mediated T-cell priming in our system, we treated B16-Ova cells with mitoxantrone, etoposide, or doxorubicin, and measured calreticulin exposure on the cell surface at 24 hours, at the time when the treated cells are exposed to BMDCs. We also analyzed HMGB1 and ATP release during the first 24 hours of chemotherapy treatment, and during the 24-48 hours post-treatment window, which corresponds to the full period of BMDC co-culture (Fig. 1A). In previous reports, etoposide was not considered an immunogenic cell death inducing drug due to its inability to induce ER stress and calreticulin exposure in CT26 cells (23), despite inducing the release of HMGB1 and ATP (31). We however included etoposide in these experiments because it induced equivalent levels of IFN-γ^+^CD8^+^ T-cells as mitoxantrone in our *in vitro* assay for DC-mediated T-cell responses. Doxorubicin was specifically chosen as the third drug for comparison because it also belongs to the same class of DNA-damaging topoisomerase II inhibitors as etoposide and mitoxantrone, but did not induce T-cell IFN-γ responses in our assay system (Fig. 1C), although it has been reported to induce calreticulin exposure and the release of HMGB1 and ATP in CT26 cells (23,31).

As shown in Figs. 3A (left panel) and S3A, using two different anti-calreticulin antibodies, all three drugs elicited only low levels of calreticulin exposure at this time point (24 hours), with <20% of the cells staining positively. Mitoxantrone-treatment induced the highest percentage of cells with externalized calreticulin when analyzed by flow cytometry after 24 hours of drug exposure. Treatment with either low or high etoposide concentrations caused an intermediate percentage of cells to display calreticulin surface exposure (<10%), while doxorubicin treatment resulted in the lowest percentage of cells with calreticulin exposure (<5%).

**Fig 3:**
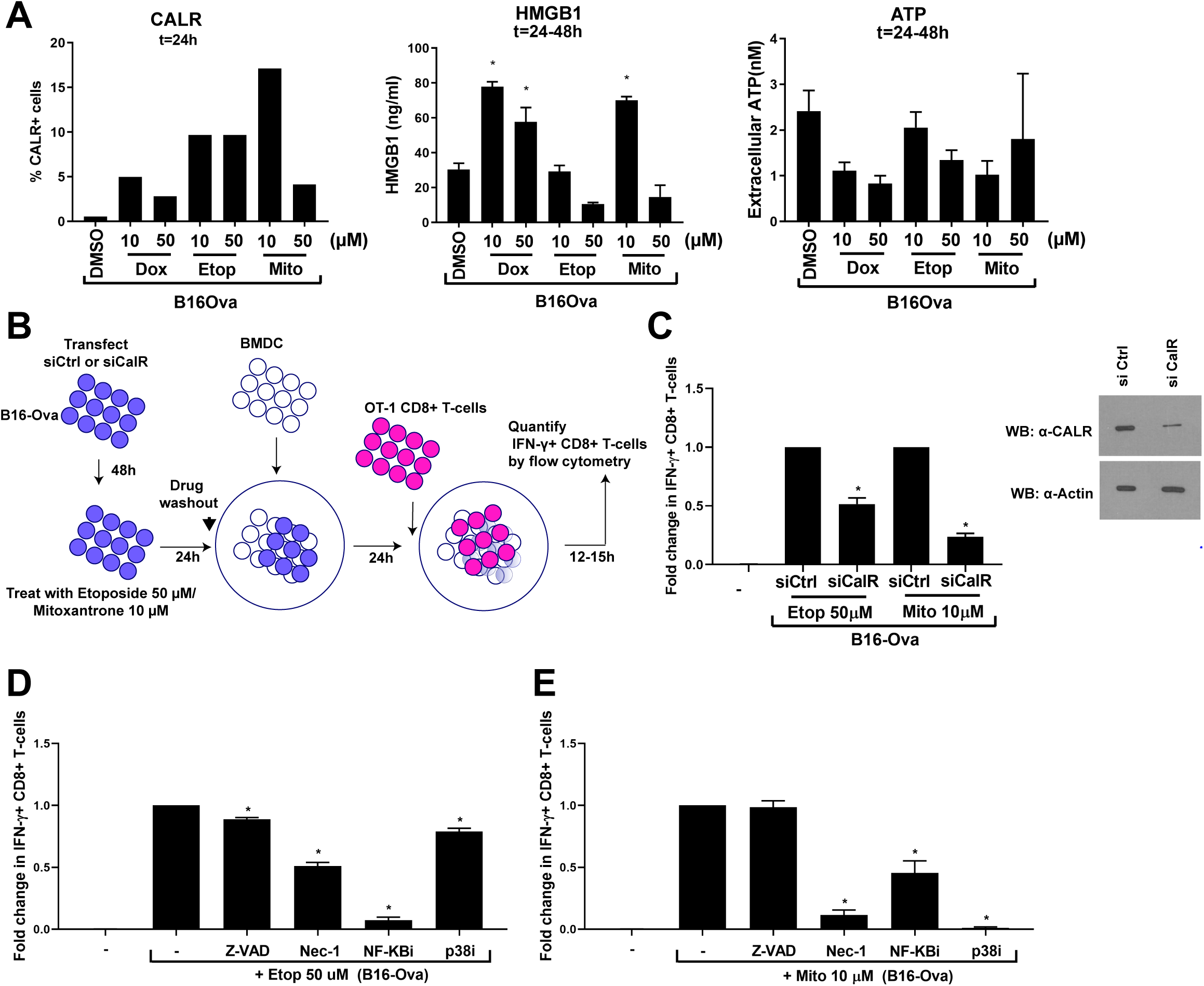
**Induction of T-cell IFN-γ by BMDC co-cultured with etoposide- or mitoxantrone-treated B16-Ova cells cannot be explained by the canonical immunogenic death markers calretilcuin, HMGB1 and ATP but is dependent on RIPK1, NF-kB and p38MAPK signaling in tumor cells**. **A**. Left panel: percentage of B16-Ova tumor cells displaying surface calreticulin 24 hours after the indicated treatment from a representative experiment. Results from an additional representative experiment using a second anti-CALR antibody is shown in Figure S3A. Middle and right panels: levels of HMGB1 and ATP in the culture media measured 24-48 hours after the indicated treatment. Results are from 4 independent experiments, with error bars indicating SEM. Data in the middle panel was analyzed by comparison to DMSO-treated controls using ANOVA followed by Dunnett’s multiple comparisons test. * indicates p<0.03. **B**. Schematic of the siRNA experiment testing the role of B16-Ova cell calreticulin in BMDC-mediated T-cell IFN-γ induction. **C**. Left panel: quantification of the fold change in %IFN-γ+ CD8+ T-cells from the experiment in panel B. Data for each calreticulin knockdown condition is normalized to the respective drug-treated control knockdown condition. Results represent 3 independent experiments with error bars indicating SEM. Data were analyzed by comparison of drug-treated calreticulin knock-down cells to their respective drug-treated control knockdown cells using a two-tailed t-test. * indicates p<0.002. Right panel: quantification of calreticulin knockdown efficiency by Western blotting. Actin was used as a loading control. First lane (-) defined as in Fig. 1C. **D and E**. Quantification of the fold change in %IFN-γ+ CD8+ T-cells induced by BMDC following incubation with etoposide- or mitoxantrone-treated B16-Ova cells that were co-treated with the indicated DNA damaging agent plus either Z-VAD, Necrostatin-1 (Nec- 1), Bay11-7085 (inhibitor of NF-kB signaling) or SB202190 (p38 MAPK inhibitor).. First lane (-) defined as in Fig. 1C. Data in D and E are normalized to the condition in which B16-Ova cells are treated with etoposide alone or mitoxantrone alone, respectively. Results represent 3 independent experiments with error bars indicating SEM. * indicates p<0.035 wherein each co-treatment condition (etoposide or mitoxantrone + either Z-VAD, Nec-1, NF-kBi or p38i) is compared with the etoposide or mitoxantrone alone condition using ANOVA followed by Dunnett’s multiple comparisons test.

As shown in Fig 3A (middle and right panels), there was no correlation between HMGB1 or ATP release and induction of immunogenicity as measured by IFN-γ production in stimulated T-cells. Cells treated with etoposide showed the lowest levels of HMGB1 release into the media during the 24-48 hours post-treatment window (i.e. when the cells are exposed to BMDCs) (Fig. 3A-middle panel), despite being highly immunogenic. In contrast, doxorubicin treatment led to high levels of HMGB1 release, similar to what was observed with mitoxantrone (10 µM) treatment, despite its inability to promote BMDC-mediated T-cell IFN-γ responses. Similar trends in HMGB1 release were observed during the first 24 hours of treatment (Fig. S1B). During the first 24 hours of drug exposure, ATP release from B16-Ova cells following doxorubicin and mitoxantrone treatment was markedly higher compared to etoposide (Fig. S3C), despite the observation that only mitoxantrone and etoposide-treated B16Ova cells induced DC-mediated IFN-γ in T-cells. Levels of ATP release following etoposide treatment of B16-Ova cells were not statistically significantly different than those levels of ATP released by DMSO-treated controls (Fig. S3C). During the 24-48 hour window after drug exposure, when B16-Ova cells are co-cultured with BMDC, the release of ATP by B16-Ova cells for all drug/dose treatment conditions was comparable to the baseline levels induced by the vehicle control condition (Fig. 3A, right panel). Overall, neither HMGB1 release nor ATP secretion were predictive of DC-mediated T-cell IFN-γ responses for all three chemotherapy agents.

Given the lack of correlation between HMGB1 or ATP release from the drug-treated tumor cells and dose-specific induction of IFNγ in T-cells, we focused on the specific role of surface calreticulin exposure. To directly measure the contribution of calreticulin to DC-mediated T-cell IFN-γ responses in our assay, the experiments outlined in Fig. 1A were repeated following siRNA knock-down of calreticulin in B16-Ova cells (Fig. 3D). As shown in Fig. 3B and C, siRNA knockdown of calreticulin prior to mitoxantrone treatment reduced the percentage of DC-mediated IFN-γ^+^ T-cells by ∼80%. In contrast, following etoposide treatment, calreticulin knock-down reduced the percentage of DC-mediated IFN-γ^+^ T-cells by only ∼50% compared to siRNA controls. These data suggest that in B16-Ova cells, calreticulin can account for the majority, but not all of the mitoxantrone-mediated immunogenicity, but is of somewhat lesser importance for BMDC-mediated T-cell IFN-γ induction in response to etoposide-treatment. In both cases additional genotoxic stress signaling mechanisms are likely to be involved..

### Induction of T-cell IFN-γ responses by BMDC co-cultured with etoposide- or mitoxantrone-treated tumor cells is dependent on RIPK1, NF-kB and p38MAPK signaling in tumor cells

To test this, we inhibited RIPK1 (a known determinant of necroptosis) (32), caspases, (known determinants of apoptosis and pyroptosis) (33), NF-kB signaling (a critical regulatory node for survival and cytokine production) (34) or p38MAPK (a well known master regulator of stress signaling, including those downstream of DNA-damage) (35). B16-Ova cells were co-treated with etoposide or mitoxantrone in combination with the RIPK1 inhibitor necrostatin-1 (Nec-1), the pan-caspase inhibitor Z-VAD, the NF-kB signaling inhibitor Bay11-7085 (36) or the p38MAPK inhibitor SB202190 (37), prior to co-culture with BMDC. As shown in Figs. 3D and E, co-treatment with Nec-1 inhibited the ability of both etoposide and mitoxantrone-treated B16-Ova cells, co-cultured with BMDCs, to induce IFN-γ in T-cells, suggesting that the ability of both etoposide and mitoxantrone to induce immunogenicity in this model is RIPK1-dependent. In contrast, co-treatment of B16-Ova cells with Z-VAD only marginally reduced T-cell IFN-γ responses (by ∼12%) with etoposide and had no effect with mitoxantrone, indicating that the process was largely independent of caspases for both agents. Furthermore, co-treatment of B16-Ova cells with the NF-kB signaling inhibitor Bay11-7085 and etoposide reduced the frequency of IFN-γ+ T-cells by >90% while co-treatment with Bay 11-7085 and mitoxantrone reduced the frequency of IFN-γ+ T-cells by >50% suggesting that NF-kB signaling in both etoposide and mitoxantrone-treated B16-Ova cells is important for the induction of DC-mediated T-cell IFN-γ responses. Finally, co-treatment of B16-Ova cells with the p38 MAPK inhibitor SB202190 and etoposide reduced the frequency of IFN-γ+ T-cells by ∼22% while co-treatment with SB202190 and mitoxantrone nearly abrogated the induction of IFN-γ+ T-cells altogether. Consistent with these results, both etoposide and mitoxantrone, which induced DC-mediated T-cell IFN-γ responses, but not doxorubicin, which did not, were found to induce RIPK1 activation in B16-Ova cells as shown by western blotting with an anti-phoshoRIPK1(S166) antibody (Fig. S3D). Furthermore, as shown in Fig S3E, western blotting of cell lysates with an anti-phospho-p38 antibody demonstrated p38MAPK activation by etoposide and mitoxantrone, as well as doxorubicin (which did not induce a DC-mediated T-cell IFN-γ response), suggesting that induction of p38MAPK signaling in tumor cells is necessary but not sufficient for the induction of IFN-γ in T-cells. Taken together, these data suggest that active signaling through the RIPK1, NF-kB and p38MAPK signaling pathways in live but damaged tumor cells following chemotherapy treatment is necessary for the induction of DC-mediated T-cell IFN-γ responses.

### In situ treatment of B16-Ova tumors in mice with etoposide does not synergize with systemic checkpoint blockade consistent with the marked reduction of DC-mediated T-cell IFN-γ responses when DCs and/or T-cells are exposed to etoposide

Given the ability of etoposide-treated B16-Ova cells to induce DC-mediated T-cell IFN-γ responses *ex vivo*, as shown in Fig. 1, we reasoned that intra-tumoral administration of etoposide could enhance DC function *in vivo* by increasing the immunogenicity of B16-Ova cells. This would be expected to induce antigen-specific T-cell expansion *in vivo*, particularly if used in combination with systemic immune checkpoint blockade. To test this, mice bearing flank B16-Ova tumors were treated by intra-tumoral administration of either saline or etoposide (three weekly doses) in the presence or absence of systemic anti-PD1 and anti-CTLA4 antibodies (two doses a week for three weeks) to confer immune checkpoint blockade (Fig. 4A). As shown in the upper panels of Fig. 4B, intra-tumoral injection of etoposide alone had no effect on tumor growth. Systemic administration of immune checkpoint blockade in combination with intra-tumoral chemotherapy also did not significantly enhance survival beyond that seen with immune checkpoint blockade alone (Figs. 4B-C). Furthermore, when we examined the frequency of circulating H2-K^b^/SIINFEKL-specific CD8^+^ T-cells, we were unable to detect an expansion of these cells when compared to the group that received checkpoint blockade alone (Fig. 4D).

**Fig 4:**
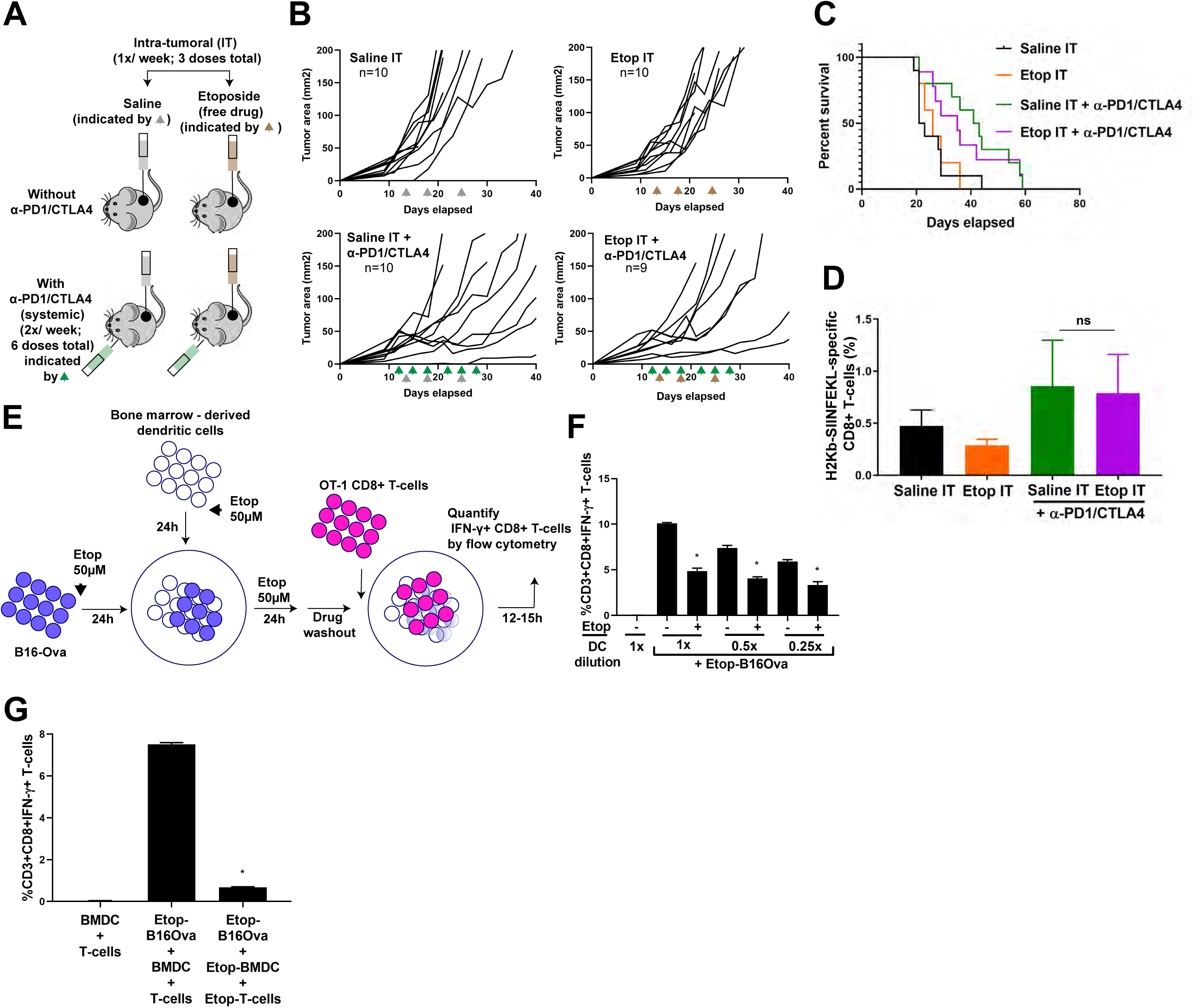
**Intra-tumoral administration of etoposide does not induce T-cell expansion *in vivo* and shows no additional therapeutic benefit compared to checkpoint blockade alone**. **A**. Schematic of the experimental design and dosing regimen used for testing intra-tumoral administration of etoposide in the presence or absence of systemic anti-PD1 and anti-CTLA4. **B**. Tumor growth curves in mice bearing B16-Ova tumors treated with intra-tumoral saline (Saline IT) (gray arrowheads) or etoposide (Etop IT) (brown arrowheads) in the presence or absence of systemic anti-PD1 and anti-CTLA4 (green arrowheads). The number of mice in each group is indicated. One mouse in the Etop IT + anti-PD1/CTLA4 group did not show tumor growth beyond 4mm^2^ throughout the experiment and was excluded. **C**. Kaplan-Meier survival curves of the experiment in panel B. Survival of the Etop IT + anti-PD1/CTLA4 treatment group was not significantly different from that of the Saline IT + anti-PD1/CTLA4 group (log-rank test). **D**. Frequency of circulating H2-K^b^/SIINFEKL -specific CD8+ T-cells from mice treated with the conditions indicated. Error bars represent SEM. The frequency of H2-K^b^/SIINFEKL tetramer stained CD8+ T-cells in the Etop IT + anti-PD1/CTLA4 treatment group was not significantly different from that of the Saline IT + anti-PD1/CTLA4 group (one-tailed t-test, p=0.4553). **E**. Schematic of the experiment examining etoposide co-treatment of BMDC and B16- Ova cells prior to the addition of T-cells. **F**. Quantification of IFN-γ+ CD8+ T-cells from the experiment in panel E. Error bars represent SEM. * indicates p<0.0001, p<0.0005, p<0.002 respectively for DC number dilutions 1x, 0.5x and 0.25x compared to their respective negative (-) controls (one-tailed t-test with Bonferroni correction). **G**. Quantification of IFN-γ+ CD8+ T-cells induced by BMDC after co-culture with etoposide-treated B16-Ova cells when both BMDC and T-cells were exposed to etoposide compared to when only B16-Ova cells were exposed. Error bars represent SEM. * indicates p<0.0001 (one-tailed t-test).

Intra-tumoral administration of etoposide, however, exposes both tumor cells and non-tumor cell types such as intra-tumoral DCs to this cytotoxic drug, which could potentially limit DC activation and impair the expansion of tumor antigen-specific T-cells. To test this hypothesis, we revised the assay shown in Fig. 1A to now include co-exposure of both the BMDCs and tumor cells to etoposide prior to the addition of OT-1 T-cells (Fig. 4E). As shown in Fig. 4F, co-exposure of both BMDCs and tumor cells to etoposide significantly reduced the appearance of IFN-γ^+^CD8^+^ T-cells compared to exposure of B16-Ova cells alone, indicating that exposure of DCs to etoposide impairs their ability to induce T-cell IFN-γ responses. Consistent with this idea, the viability of BMDCs was significantly reduced upon exposure to etoposide (Fig. S4). We further modified the assay to include exposure of all of the relevant cell types - tumor cells, BMDCs and T-cells - to etoposide, mirroring what might occur following intra-tumoral injection of the drug *in vivo*. As shown in Fig. 4G, this triple co-exposure resulted in an even more profound loss of IFN-γ+ T-cells to less than 10% of the level seen when etoposide exposure is limited to the tumor cells alone.

### Intra-tumoral injection of ex vivo etoposide-treated tumor cells synergizes with immune checkpoint blockade, enhances survival and induces resistance to re-challenge

Exposure of BMDC and T-cells to etoposide reduced the induction of IFN-γ^+^CD8^+^ T-cells by drug-treated B16-Ova cells compared to etoposide exposure of B16-Ova cells alone. We therefore reasoned that the intra-tumoral injection of *ex vivo* etoposide-treated B16-Ova cells into B16-Ova tumors *in vivo*, rather than intra-tumoral injection of the free drug, would minimize exposure of other immune cell types in the tumor and draining lymph node to the cytotoxic effects of etoposide. To test this, mice bearing flank B16-Ova tumors received intra-tumoral injection of either saline or *ex vivo* etoposide-treated B16-Ova cells in the presence or absence of systemic checkpoint blockade (Fig. 5A). As shown in Figs. 5B-D, intra-tumoral administration of *ex vivo* etoposide-treated tumor cells alone had no effect on subsequent tumor progression. However, when used in combination with systemic checkpoint blockade, the mice displayed superior tumor control compared to those that received checkpoint blockade alone, resulting in complete tumor regressions in ∼35% of mice. Furthermore, survival was also markedly enhanced in this group (Fig. 5C). Importantly, analysis of circulating lymphocytes in these animals revealed an enhanced frequency of H2-K^b^/SIINFEKL-specific CD8+ T-cells (Fig. 5E), indicating that intra-tumoral administration of *ex vivo* etoposide-treated tumor cells functions as an effective injured cell adjuvant, which in combination with immune checkpoint blockade, promotes efficient T-cell priming and anti-tumor immunity. The subset of mice that demonstrated complete tumor regression after injured cell adjuvant treatment remained tumor-free for at least 98 days (Fig. 5C). These complete responders and naive control mice (which were never previously exposed to B16-Ova tumor cells) were re-challenged in the contralateral flank with live B16-Ova cells (Fig. 5F, left panel). As shown in the right panel of Fig. 5F, tumors grew to 200 mm^2^ cross-sectional area within 30 days in the naive mice, (at which point they were euthanized). Notably, none of the intra-tumoral vaccine-treated animals who were cured of their initial tumors after therapy developed tumors upon re-challenge, suggesting that combining systemic checkpoint blockade with intra-tumoral injections of the injured cell adjuvant induces anti-tumor immunological memory.

**Fig 5:**
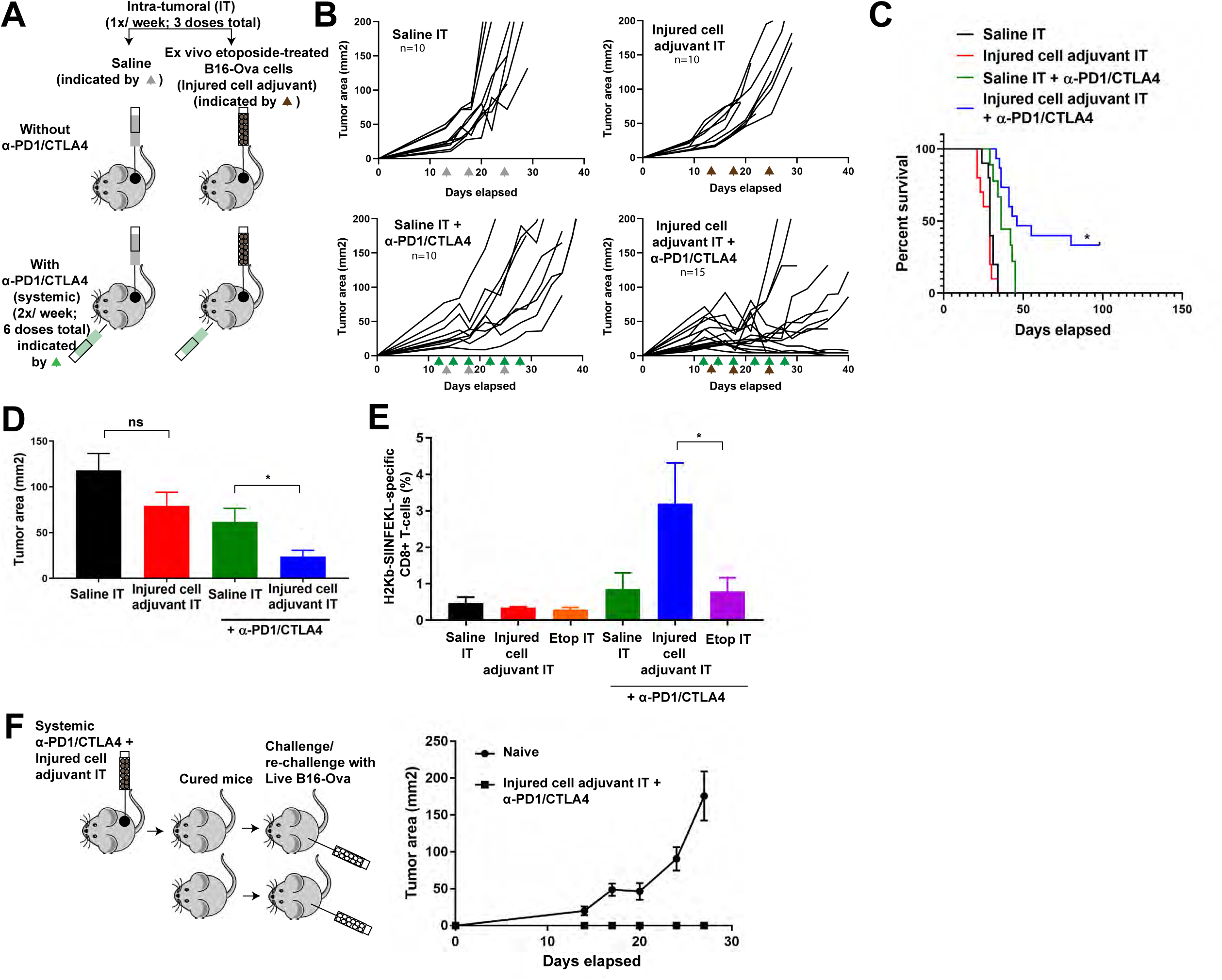
**Intra-tumoral administration of *ex vivo* etoposide-treated B16-Ova cells (injured cell adjuvant) in combination with systemic checkpoint blockade increases anti-tumor CD8+ T-cell expansion, shows enhanced therapeutic benefit and induces long-term immunological memory**. **A**. Schematic of the experimental design and dosing regimen used for testing intra-tumoral administration of etoposide-treated B16-Ova cells (injured cell adjuvant) in the presence or absence of systemic anti-PD1 and anti-CTLA4. **B**. Tumor growth curves for mice treated with intra-tumoral saline (Saline IT) (gray arrowheads) or *ex vivo* etoposide-treated B16-Ova cells (Injured cell adjuvant IT) (brown arrowheads) in the presence or absence of systemic anti-PD1 and anti-CTLA4 (green arrowheads). ‘n’ indicates the number of mice in each group. **C**. Kaplan-Meier Survival curves of the experiment in panel B. * indicates p<0.02 when compared to the group treated with Saline IT + anti-PD1/CTLA4 (log-rank test). **D**. Average tumor cross-sectional area on Day 21 for each treatment group. Error bars indicate SEM. * indicates p<0.02 when compared to the group treated with Saline IT + anti-PD1/CTLA4 (one-tailed t-test). **E**. Frequency of circulating H2-K^b^/SIINFEKL -specific CD8+ T-cells from mice following the indicated treatments. Treatment groups shown in Fig 4D are also included for comparison. * indicates p<0.04 (one-tailed t-test). **F**. Left panel: Schematic of the experiment in which 5 naïve mice and 5 mice that demonstrated complete tumor regression following treatment with injured cell adjuvant + systemic anti-PD1/CTLA4 were re-challenged in the opposite flank with 100,000 live B16- Ova cells. Right panel: Resulting tumor growth curves. Error bars indicate SEM.

To examine whether this response was unique to the B16 cell line, or to cells engineered to express the ovalbumin antigen, we performed similar intra-tumoral injections of saline- or etoposide-treated tumor cells, in the presence or absence of systemic immune checkpoint blockade, with MC-38 murine colon carcinoma cells that do not express ovalbumin, (Fig. S5A). Figs. S5A and B show that in this tumor model there was minimal benefit of immune checkpoint blockade alone when the MC-38 tumors were injected with saline. Similarly, intra-tumoral injection of etoposide-treated MC-38 tumor cells into pre-existing MC-38 tumors failed to elicit an anti-tumor immune response in the absence of systemic immune checkpoint blockade. However, 20% of the animals who received the combination of the MC-38 tumor cell vaccine together with systemic immune checkpoint blockade showed complete tumor regression and prolonged survival.

### Batf3 (-/-) mice do not respond to the injured cell adjuvant and checkpoint blockade combination

To test whether the efficacy of the injured cell adjuvant in combination with immune checkpoint blockade treatment for an anti-tumor immune response depends on DCs that can cross-present tumor antigens, we enumerated the numbers of CD11b^+^C103^-^ DC2 cells and CD11b^-^CD103^+^ DC1 cells by immunophenotyping and flow cytometry (Fig. S6). CD11b-CD103+ DC1 cells, which are typically also Batf3+ (38,39), are known to cross present tumor antigens to CD8+ T-cells (40). As before, mice bearing flank B16-Ova tumors were treated with saline or the injured cell adjuvant intra-tumorally, in the presence or absence of systemic checkpoint blockade (Fig. 6A), and analyzed. In addition, we also included a cohort that were treated with intra-tumoral etoposide in combination with checkpoint blockade. After 2 doses of the injured cell adjuvant or etoposide and 3 doses of checkpoint blockade, immunophenotyping of the tumors revealed an enhanced number of CD103^+^ DC1 in tumors that were being treated with the injured cell adjuvant and checkpoint blockade, compared to the other groups (Fig 6B). In addition, cross-sections of tumors treated with the injured cell adjuvant and checkpoint blockade showed markedly enhanced Batf3 staining by immunohistochemistry (Fig. 6C) indicating the enhanced presence of Batf3^+^ DC, which was not present in the other treatment groups. Intra-tumoral injection of free etoposide combined with checkpoint blockade did not enhance numbers of CD103^+^ DC1, consistent with the lack of T-cell expansion and lack of efficacy seen *in vivo* (Fig. 5A-D) and *in vitro* (Fig. 4E-G) with this treatment.

**Fig 6:**
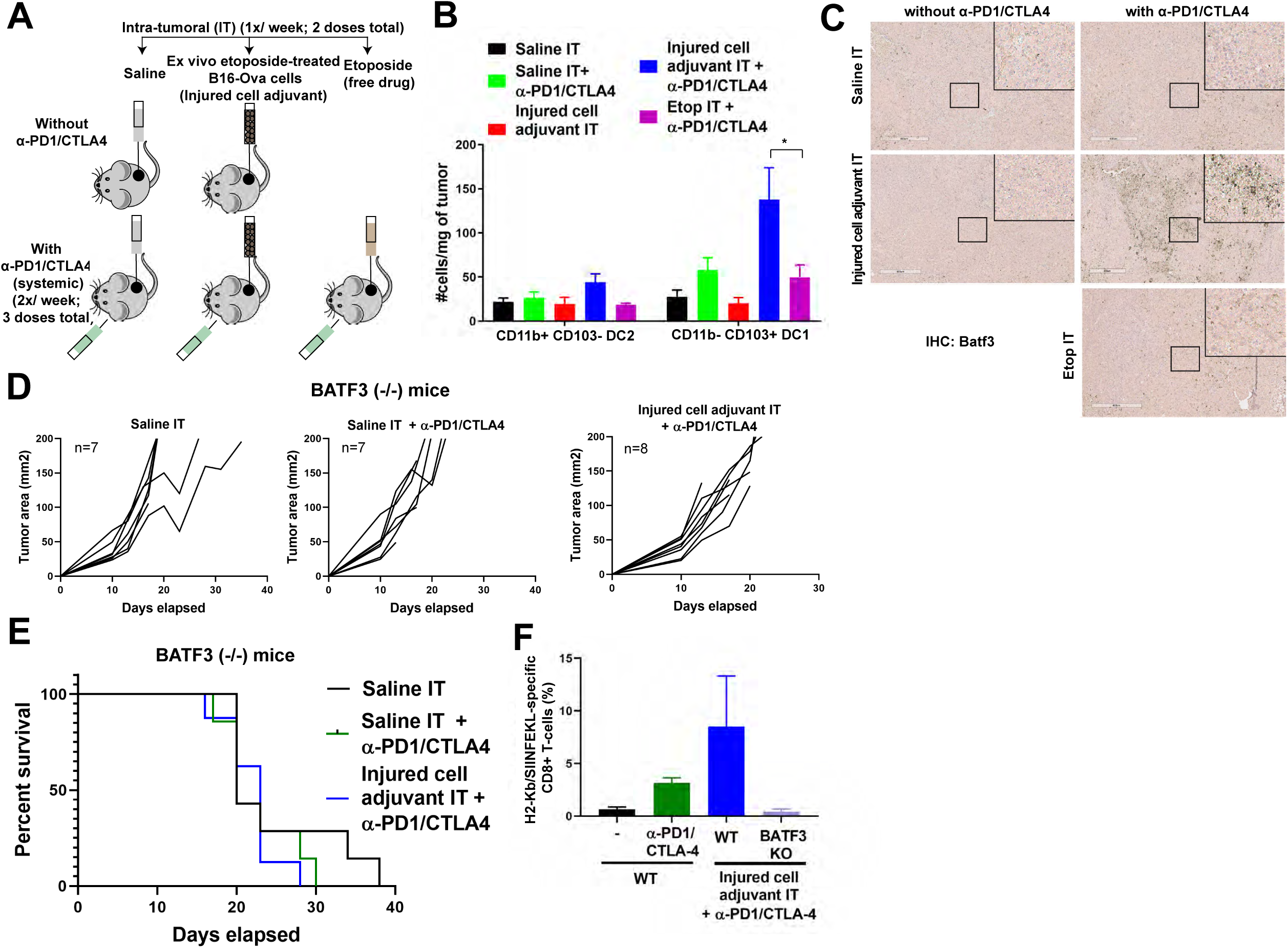
**Intra-tumoral administration of the injured cell adjuvant in combination with systemic checkpoint blockade increases the numbers of CD11b-CD103+ DC (DC1) in the tumor and is not effective therapeutically in Batf3 deficient mice**. **A**. Schematic of the experimental design and dosing regimen used to test the effect of intra-tumoral etoposide-treated B16-Ova cells in combination with systemic anti- PD1/CTLA4, on the frequency of intra-tumoral DC. **B**. Quantification of intra-tumoral CD11b-CD103+ DC1 and CD11b+CD103- DC2 subsets from treated tumors analyzed by flow cytometry. Error bars represent SEM. * indicates p<0.04 (one-tailed t-test). **C**. Tumor sections were stained with an anti-Batf3 antibody. Insets show higher magnification images of the boxed central region of each section. Scale bar indicates 400 μm. **D**. Tumor growth curves of Batf3(-/-) mice treated with intra-tumoral saline or etoposide-treated B16-Ova cells (injured cell adjuvant) in combination with systemic anti-PD1 and anti-CTLA4 antibodies. ‘n’ indicates the number of mice in each group. **E**. Kaplan-Meier survival curves of the experiment shown in panel D. The survival curves are not significantly different (log-rank test, p=0.5220). **F**. Frequency of circulating H2-K^b^/SIINFEKL specific CD8+ T-cells from WT and BATF3 (-/-) mice treated with the conditions indicated.

To directly validate the contribution of Batf3^+^CD11b^-^CD103^+^ DC1 cells to anti-tumor immunity induced by the combination of our injured cell adjuvant and checkpoint blockade, the experiment shown in Fig. 5A was repeated using Batf3^-/-^ mice. As shown in Figs. 6D and E, intra-tumoral injection of *ex vivo* etoposide-treated tumor cells with systemic immune checkpoint blockade failed to induce tumor control or prolong the lifespan of tumor-bearing mice in the absence of Batf3. Lastly, while the injured cell adjuvant and systemic ICI combination enhanced the frequency of circulating H2-K^b^/SIINFEKL-reactive CD8^+^ T-cells in WT mice, there was no such increase in Batf3-deficient mice (Fig. 6F). Taken together, these data strongly suggest that intra-tumoral administration of *ex vivo* etoposide-treated tumor cells as an injured cell adjuvant, in combination with systemic checkpoint blockade, promotes Batf3^+^ DC-mediated anti-tumor T-cell responses leading to improved survival, and complete tumor regressions in a subset of mice concurrent with long-term anti-tumor immunological memory.

## DISCUSSION

The use of conventional DNA-damaging chemotherapy to induce tumor cell stress and/or death and thereby further augment immunogenicity is an attractive idea (26). However, there has been no systematic way to discover and achieve synergy by combining chemotherapy with ICI in present clinical practice. In this study, we used an *in vitro* experimental system to identify specific chemotherapeutic drug/dose combinations to treat tumor cells wherein the treated tumor cells, co-cultured with DC, enhance IFN-γ induction in T-cells. However, if DCs and/or T-cells are also exposed to the chemotherapy drug, T-cell IFN-γ responses are impaired and consistently, direct intra-tumoral injection of chemotherapy as a free drug, in combination with systemic ICI administration, was largely ineffective. Dead cells or cell-free supernatants generated after chemotherapy treatment, when co-incubated with BMDC, were not sufficient to promote T-cell IFN-γ responses. Notably, we show that active signaling through RIPK1. NF-kB and p38MAPK pathways in live injured cells is necessary for T-cell activation following DC exposure to either mitoxantrone- or etoposide-treated tumor cells. We have further identified that *ex vivo* chemotherapy-treated tumor cells, function as an injured cell adjuvant, when administered intra-tumorally, in combination with systemic ICI in mouse cancer models (Fig 7). Using this combination, we observed an expansion of CD103^+^ intra-tumoral DCs, an increase in the frequency of H2-K^b^/SIINFEKL-reactive circulating anti-tumor CD8^+^ T-cells, and markedly enhanced tumor control and significant survival benefit compared to ICI alone. Furthermore, a subset of mice showed complete tumor regressions and resistance to re-challenge with live tumor cells in the contra-lateral flank. A similar response was observed using MC-38 cells lacking ovalbumin, indicating that the results were not limited to one tumor cell type, or to cells that express a foreign non-tumor antigen.

**Fig 7:**
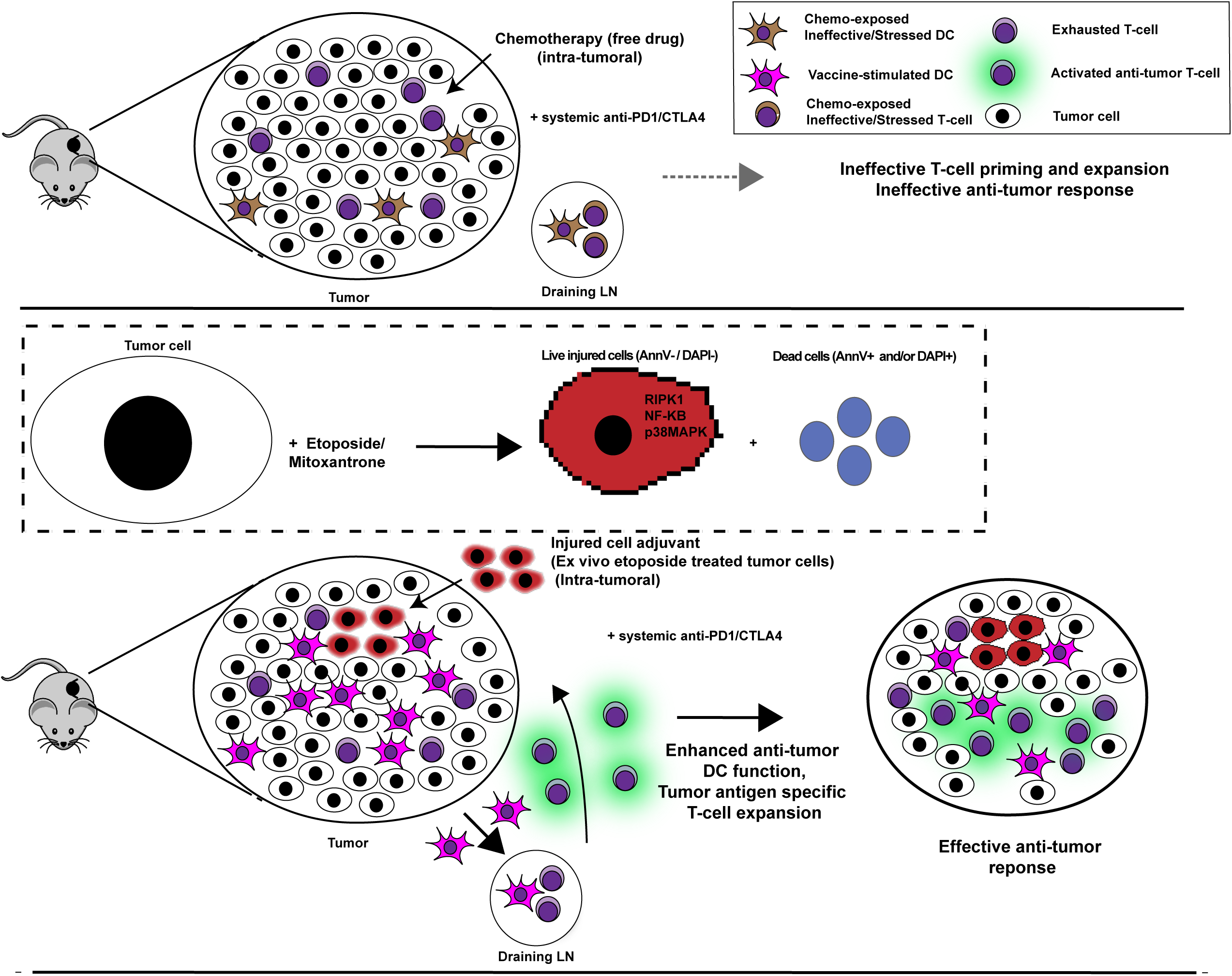
**Model of therapeutic efficacy resulting from intra-tumoral administration of *ex vivo* chemotherapy-treated tumor cells in combination with systemic immune checkpoint blockade**. Intra-tumoral injection of ex-vivo DNA damaging chemnotherapy-treated tumor cells promotes effective DC-mediated T-cell priming and expansion when combined with systemic ICI, while intra-tumoral injection of free cytotoxic drug is ineffective.

Our finding that certain types of DNA-damaging chemotherapy could increase the immunogenicity of the treated tumor cells is in good agreement with many findings from Obeid et al (23), but contrasts with other findings by the same authors. Similar to what those authors described as markers of immunogenicity, we found that *in vitro* treatment of tumor cells with mitoxantrone induced strong DC-dependent T-cell priming that could largely be explained by calreticulin exposure. Treatment of tumor cells with etoposide resulted in lower levels of exposure of calreticulin, again in agreement with Obeid et al. who reported that this agent did not induce strong immunogenic cell death. However, in contrast to that prior work, in our *in vitro* assay for T-cell priming, etoposide performed just as well as mitoxantrone. Importantly, the immunogenicity assay used by Obeid et al differs substantially from the assay we used. In their system, drug-treated tumor cells were injected into the flank of naive mice, and the mice then challenged with undamaged tumor cells injected into the opposite flank 7 days later. Failure of the second tumor cell challenge to establish a tumor was taken as evidence of anti-tumor immunity. In our system, we directly measured the ability of drug-treated cells to drive IFN-γ production in CD8+ T-cells, and further validated this effect *in vivo* for etoposide treatment by injection of the drug-treated tumor cells into pre-existing mouse tumors, followed by direct measurements of tumor response and tumor-infiltrating immune cells in the presence or absence of systemic immune checkpoint inhibitors.

In our experiments, knock-down of calreticulin prior to etoposide exposure only partially reduced the ability of these cells to induce DC-dependent T-cell priming, which could also not be explained by drug-induced HMGB1 or ATP release. Together with our finding that the dead cells or cell-free supernatants alone, or in combination, when co-incubated with BMDC, were not sufficient to induce IFN-γ in T-cells and that active signaling in the live injured fraction of cells after etoposide or mitoxantrone treatment is necessary for DC-mediated T-cell IFN-γ responses raises several interesting possibilities about the mechanisms involved in promoting effective cross-presentation of tumor antigens by DCs to T-cells. Current understanding presumes that a property of dead cells generated by chemotherapy, such as specific molecules presented on the cell surface or released into the microenvironment, are the major determinants of effective cross-presentation of tumor antigens by DC to T-cells. Our findings suggest instead that active signaling through RIPK1, NF-kB and p38MAPK by live but stressed and injured cells after chemotherapy treatment are a major determinant of efficient DC-mediated T-cell priming. However, our results do not exclude a contribution from chemotherapy-induced cell death, since some of the live injured cells after chemotherapy treatment may die during the co-incubation period with BMDCs. Finally, lysates of the chemotherapy-treated cell mixture generated by three cycles of freeze-thawing, when co-incubated with DC, do not promote T-cell IFN-γ response suggesting that an active cellular process beyond cytokine secretion may be involved. Elucidating the molecular basis of this effect will be the subject of future studies.

Our finding of RIPK1 and NF-kB involvement in driving immunogenic cell death following treatment of tumor cells with specific DNA damaging chemotherapeutic drugs is in excellent agreement with the recent results of Yatim et al., (41) and Snyder et al., (42). Yatim et al., found that artificial induction of RIPK3 in NIH-3T3 cells, followed by intradermal injection, induced priming of CD8^+^ T-cells *in vivo* in a DC-dependent manner that required RIPK1 activity in the RIPK3-induced cells. In addition, NF-kB activity was also required. As in our study, these authors also found that classic markers of immunogenic cell death (calreticulin, ATP, HMGB1) were insufficient to explain this DC-mediated T-cell priming and expansion. Snyder et al., reported that overexpression of a synthetic RIPK3 dimerization construct in tumor cells *in vitro*, followed by intra-tumoral administration of these cells conferred significant tumor control and anti-tumor immunity. Remarkably, transfection with a RIPK3 variant that is unable to activate RIPK1, but is still able to cause necroptotic cell death, failed to confer CD8 T-cell expansion or tumor control, demonstrating the importance of RIPK1 in this process.

Our results are in good agreement with recent studies wherein a subset of intra-tumoral dendritic cells, characterized by their surface expression of CD103 in mice and BDCA-3 in humans, was identified as having unique capabilities of cross-presenting tumor-associated antigens to CD8^+^ T-cells and recruiting T-cells to the tumor microenvironment through CXCL9/10 (40,43,44). The levels of these DCs in the tumor microenvironment was shown to correlate with better overall survival in melanoma patients receiving immune checkpoint inhibitors (45), consistent with the importance of these cells in enhancing anti-tumor immune responses.

Finally, our results suggest a potential method to translate these findings into clinical use, without requiring the need to genetically manipulate the cells to artificially drive RIPK3 dimerization. Tumor cells derived from patient tumor biopsies could be expanded and used to screen the immunogenicity of chemotherapeutic compounds to identify the optimal compound for a particular tumor using primary patient-derived or allogeneic DC and CD8^+^ T-cells. Matched tumor cells treated with the optimal compound identified could then be re-injected into the same tumor in combination with systemic checkpoint blockade. While clinical trials will be required to test efficacy, this approach has potential for patients whose cancers are accessible for intra-tumoral delivery and in whom conventional treatment options have failed and initial or acquired resistance to ICI has been observed.

## ACKNOWLEDGEMENTS

We thank Yi Wen Kong, Lucia Suarez-Lopez and Jesse Patterson for many helpful discussions and the Koch Institute’s Robert A. Swanson (1969) Biotechnology Center for technical support, including the Flow Cytometry and the Hope Babette Tang (1983) Histology core facilities.

## Supplemental Figure legends

**Fig S1:**
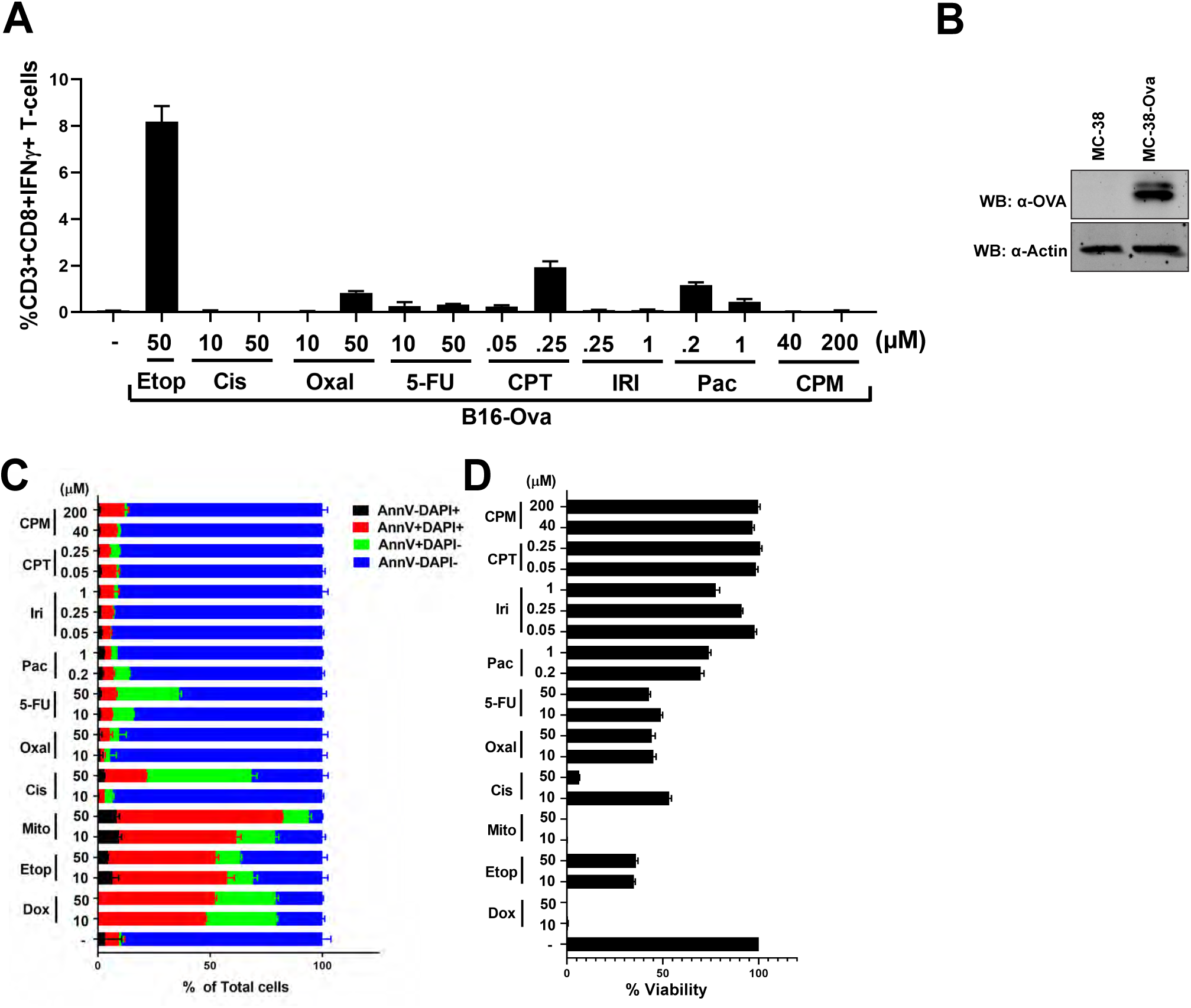
**(Related to Fig 1). Etoposide- or mitoxantrone-treated B16-Ova cells, when co-cultured with BMDC, are the most effective at inducing IFN-γ in OT-1 CD8+ T-cells**. **A**. AnnexinV/DAPI staining 48 hours after treatment with the indicated drugs and concentrations. Dox – doxorubucin, Etop – etoposide, Mito – mitoxantrone, Cis – cisplatin, Oxal – oxaliplatin, 5-FU – 5-fluorouracil, Pac – paclitaxel, Iri – irinotecan, CPT – camptothecin, CPM – cyclophosphamide. Error bars represent range obtained from at least two independent experiments. **B**. Cell viability as assessed by CellTiter-Glo signal at 48 hours after treatment with the indicated drugs and concentrations. Data is from 5 independent experiments. Error bars represent SEM. **C**. Quantification of BMDC-mediated induction of IFN-γ+ CD8+ T-cells by chemotherapy-treated B16-Ova cells. The first lane (-) indicates the percentage of IFNγ+ CD8+ T-cells produced by co-culture of BMDCs and T-cells in the absence of any B16-Ova cells. Error bars represent SEM. * indicates p< 0.006 when compared to (-) sample using ANOVA followed by Dunnett’s multiple comparisons test. **D**. Western blot of Ova expression in the MC-38-Ova cells.

**Fig S2:**
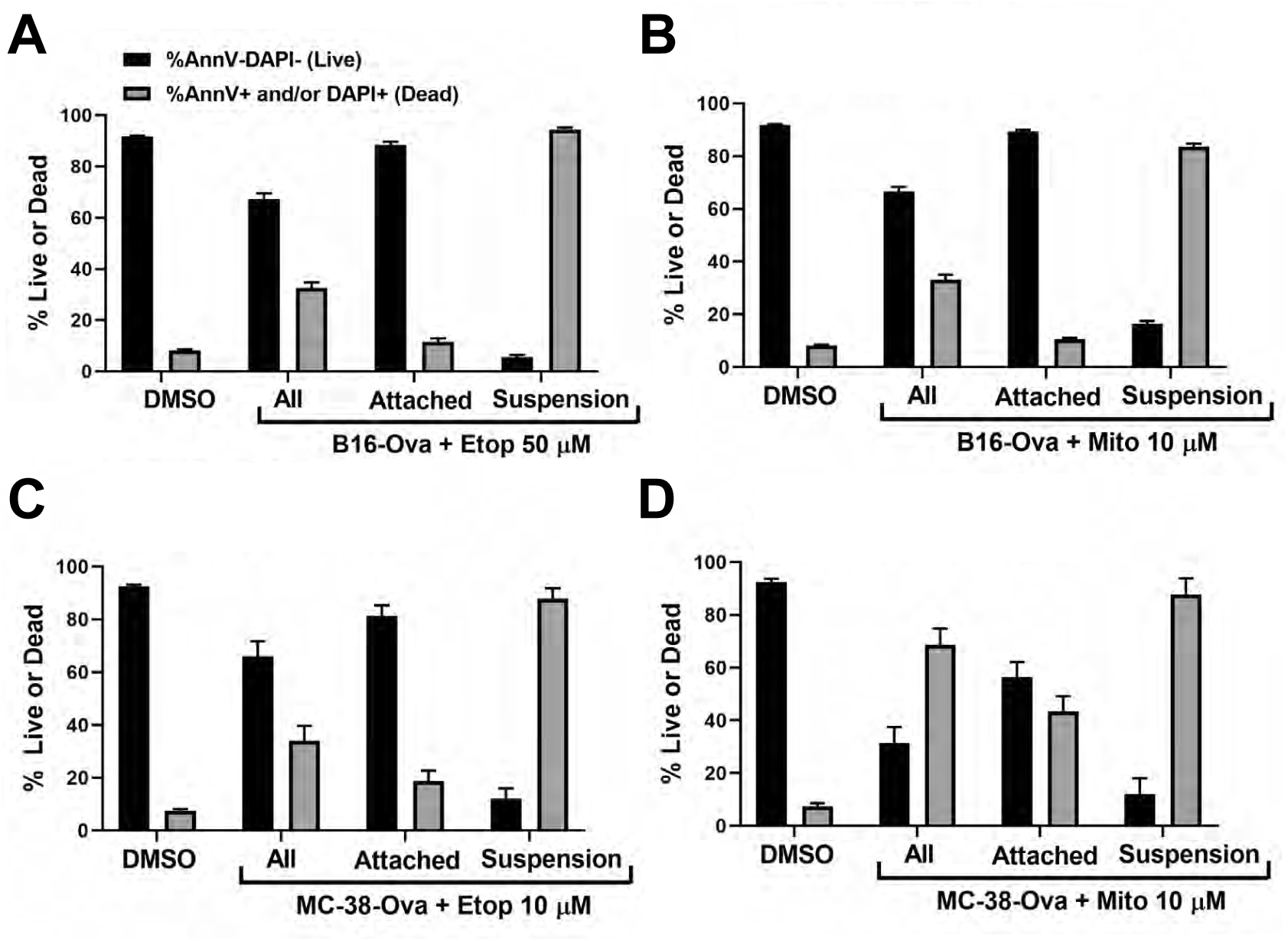
**(related to Figure 2) The attached fraction of B16-Ova and MC-38-Ova cells after etoposide or mitoxantrone treatment is predominantly live and stains AnnV and DAPI double negative while the floating (suspension) fraction is predominantly dead and stains AnnV and/or DAPI single or double positive** **A and B**. AnnV/DAPI staining as analyzed by flow cytometry of the total (all), attached, or floating (suspension) fractions of B16-Ova cells after treatment with Etoposide (50 uM) (in A) or Mitoxantrone (10 uM) (in B) for 24 hours. Quantification of live cells (AnnV and DAPI double negative; black bars) and dead cells (AnnV or DAPI single or double positive; gray bars) in each fraction from three independent experiments is shown. Errors represent SEM. **C and D**. AnnV/DAPI staining as analyzed by flow cytometry of the total (all), attached, or floating (suspension) fractions of MC-38-Ova cells after treatment with Etoposide (50 uM) (in C) or Mitoxantrone (10 uM) (in D) for 24 hours. Quantification of live cells (AnnV and DAPI double negative; black bars) and dead cells (AnnV or DAPI single or double positive; gray bars) in each fraction from three independent experiments is shown. Errors represent SEM.

**Fig S3:**
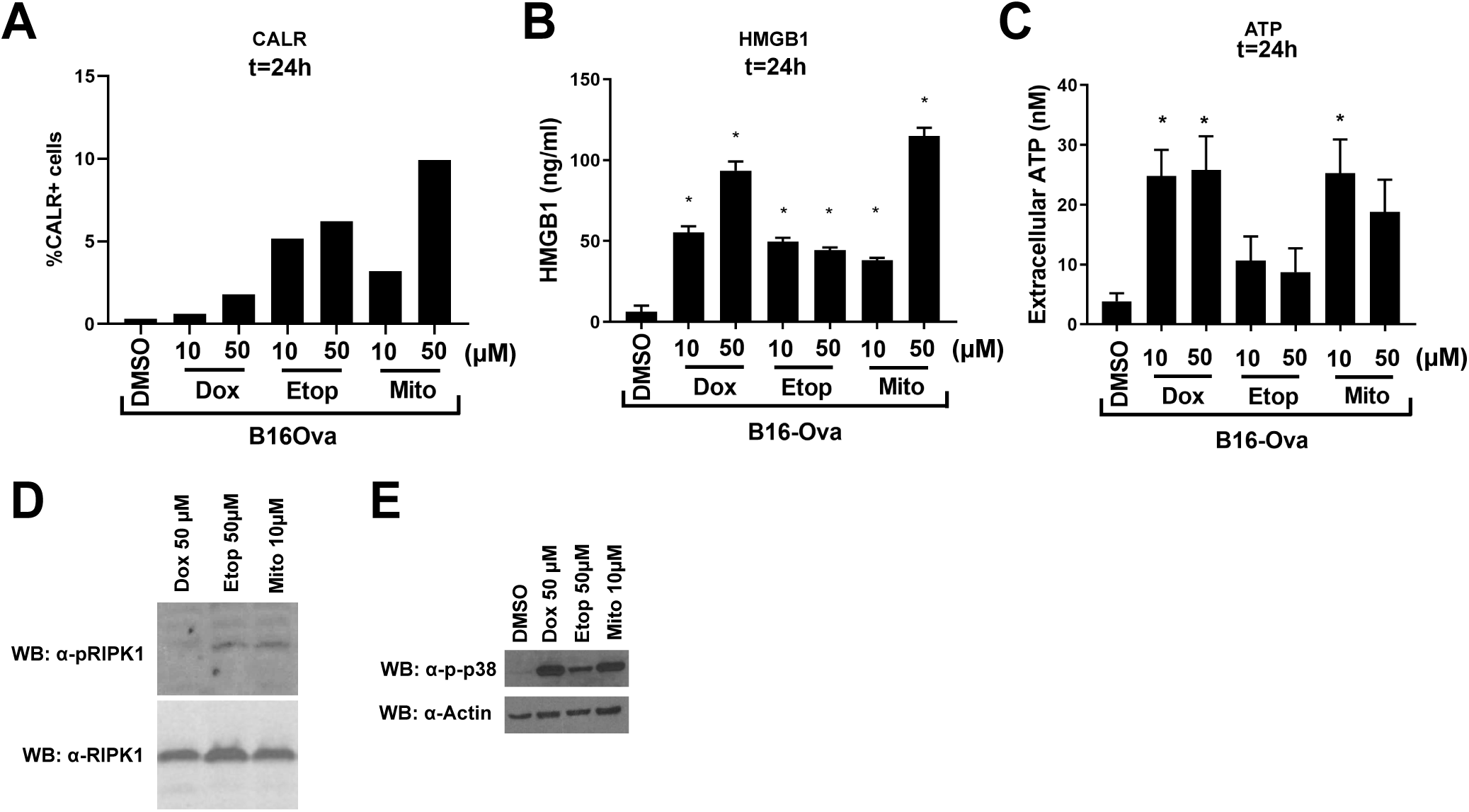
**(related to Fig 3). Levels of calreticulin exposure, HMGB1 and ATP secretion 24 hours after treatment and evidence of RIPK1 and p38MAPK activation after treatment of B16-Ova cells with doxorubicin, etoposide or mitoxantrone**. **A**. A representative experiment using an anti-CALR antibody (from CST) showing the percentage of CALR+ B16-Ova cells 24 hours after the indicated treatment. Staining with a Ctrl IgG was used to gate out background. **B**. Quantification of HMGB1 by ELISA in the cell culture media 24 hours after treatment of B16-Ova cells with the indicated chemotherapy drugs and doses. Error bars represent SEM. * indicates p< 0.0001 when compared to DMSO-treated control using ANOVA followed by Dunnett’s multiple comparisons test. **C**. Quantification of ATP by CellTiter-Glo in the cell culture media 24 hours after treatment of B16-Ova cells with the indicated chemotherapy drugs and doses. Error bars represent SEM. * indicates p< 0.027 when compared to DMSO-treated control using ANOVA followed by Dunnett’s multiple comparisons test. D. Western blotting for phospho-RIPK1 (S166) and total RIPK1 in lysates of B16-Ova cells treated with the indicated DNA damaging drugs for 15 hours. E. Western blotting for phospho-p38 and Beta-Actin in lysates of B16-Ova cells treated with the indicated DNA damaging drugs or DMSO for 15 hours.

**Fig S4:**
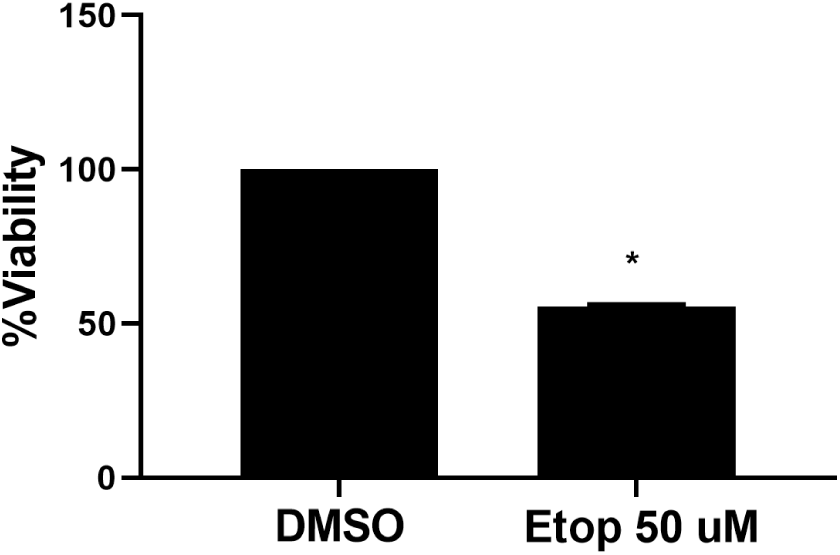
**(related to Fig 4). Exposure of BMDC to etoposide reduces viability**. BMDC viability as assessed by CellTiter-Glo at 48 hours after treatment with DMSO or etoposide. Data is from 5 independent experiments. Error bars represent SEM. * indicates p<0.0001 when compared to DMSO-treated control (one tailed t-test).

**Fig S5:**
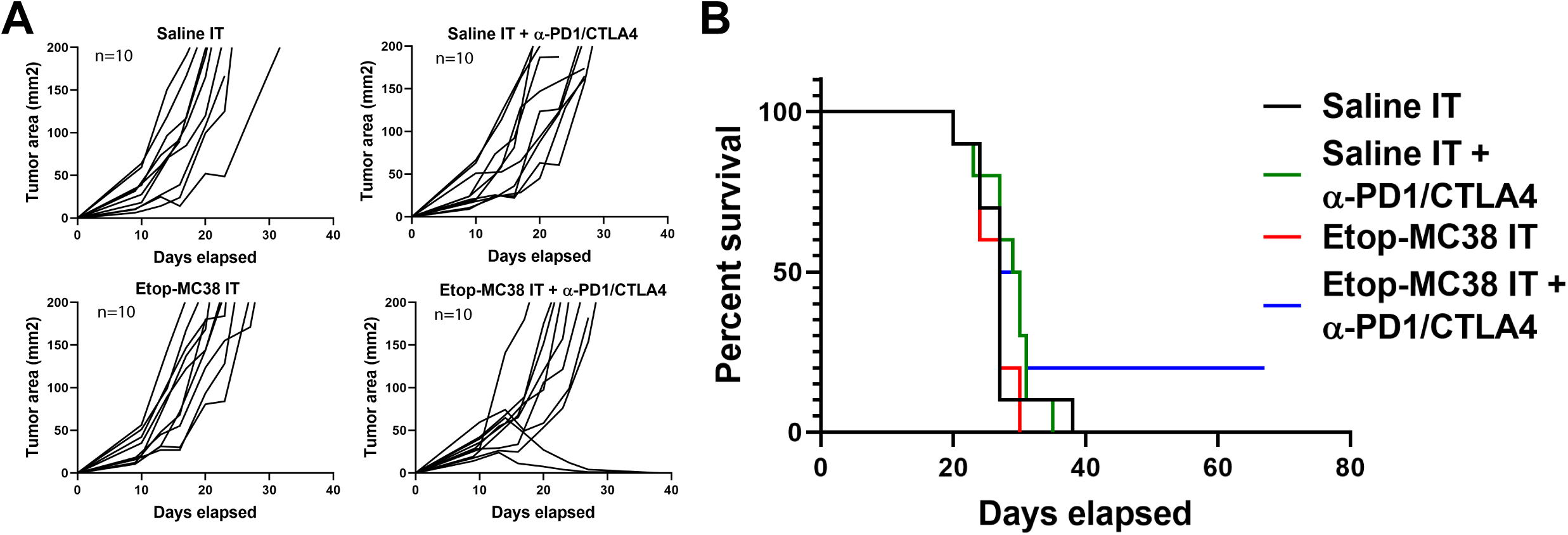
**(related to Fig 5). Intra-tumoral administration of ex vivo etoposide-treated MC-38 cells in combination with systemic checkpoint blockade shows enhanced therapeutic benefit and extended survival**. **A**. Tumor growth curves in mice bearing MC38 flank tumors treated with intra-tumoral saline (Saline IT) or *ex vivo* etoposide-treated MC38 cells in the presence or absence of systemic anti-PD1 and anti-CTLA4. ‘n’ indicates the number of mice in each group. **B**. Kaplan-Meier Survival curves of the experiment in A.

**Fig S6:**
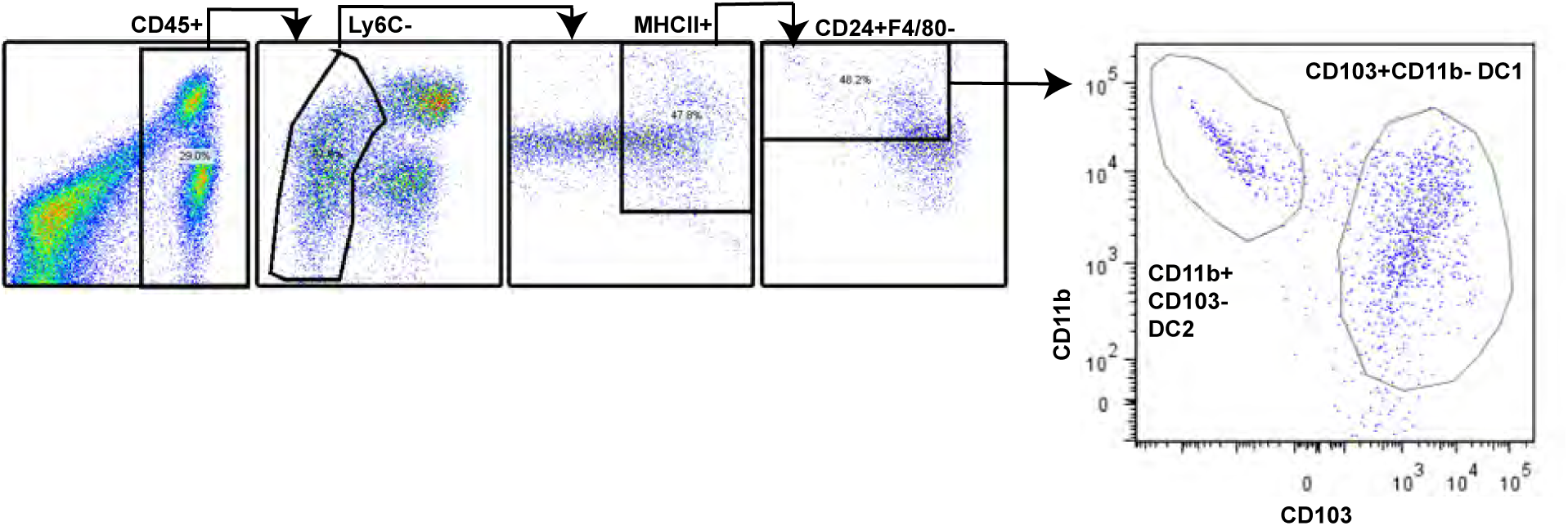
**(related to Fig 6). Gating strategy used to identify CD11b-CD103+ DC1 and CD11b+ CD103-DC2 by flow cytometry.** DCs were gated as CD45+Ly6C-MHCII+CD24+F4/80- and then further gated into DC1 (CD103+CD11b-) and DC2 (CD11b+CD103-).

